# Continuous Measures of Decision-Difficulty Captured Remotely: I. Mouse-tracking sensitivity extends to tablets and smartphones

**DOI:** 10.1101/2023.06.06.543796

**Authors:** Alexandra A. Ouellette Zuk, Jennifer K. Bertrand, Craig S. Chapman

## Abstract

As decisions require actions to have an effect on the world, measures derived from movements such as using a mouse to control a cursor on a screen provide powerful and dynamic indices of decision-making. In this first of a set of two studies, we replicated classic reach-decision paradigms across computers, tablets, and smartphones, we show that portable touch-devices can sensitively capture decision-difficulty. We see this in pre- and during-movement temporal and motoric measures across diverse decision domains. We found touchscreen interactions to more sensitively reflect decision-difficulty during movement compared to computer interactions, and the latter to be more sensitive before movement initiation. Paired with additional evidence for the flexibility and unique utility of pre- and during-movement measures, this substantiates the use of widely available touch-devices to massively extend the reach of decision science. We build upon this in the second study in this series (Bertrand et al., 2023) with the use of webcam eye-tracking to further elucidate, earlier in time, the decision process. This subsequent work provides additional support for tools that enable remote collection of rich decision data in ecologically-valid environments.

## 1 Introduction

Our lives unfold as an amalgamation of decisions made and actions taken to execute them. Ultimately, these enacted choices shape our lives and our societies. As a result, the study of human decision behaviour has inspired researchers for centuries, from interest in risk preference amongst gamblers [5], to willingness to pay given prior value contexts [27].

Historically, most measures of decision-making use verbal reports (e.g., [38, 27]), observed choices (e.g., [34]), or discrete measurements of behaviour such as reaction time and accuracy (see [41] for review). Reaction times, specifically, have been shown to reflect cognitive conflict during decision-making, with more difficult decisions leading to longer reaction times [32, 36, 40]. These approaches, which focus almost exclusively on the outcome of a decision, fail to account for the embodied nature of real-world decision-making. In the real-world, a decision is not made until a body physically enacts the choice. Recognizing that *how* we decide is likely as important as *what* we decide, researchers have started recording the dynamics of behaviour [9, 22, 14, 15, 50]. Requiring and tracking movement to select between choices, reach-decision paradigms are a popular method for continuously measuring the factors that underlie and bias the decision process. These tasks have quantified decision behaviours across a variety of choice domains for both real 3-D reaching [8, 7, 21, 22] and for 2-D computer-mouse tracking [20, 44, 26, 45].

Computerized reach-decision tasks, with 2-D movements made by a computer-mouse are a particularly sensitive, flexible, and scalable technique for the examination of decision processes ([28, 33, 26, 17, 20, 44, 31] and many more). Requiring participants to start with their mouse cursor centered at the bottom of the computer screen and necessitating the selection of one of two (most commonly) choice options located in the top left or right corners of the screen, classic mouse-tracking paradigms record the attraction toward each of the two choice options. This generates a vertical movement component relatively independent of the competition between options (though, movement speed has been related to different aspects of the decision process [14, 15]) and a critical horizontal movement component that tracks either directly toward one of the two options when there is no choice-competition, or indirectly between the two options when the choice-competition is high [14, 15, 44]. The typical result is a continuum of direct to indirect trajectories, reflecting the strength of competition between choice options and thus the relative difficulty of the decision. Metrics quantifying relative reach directness include the maximum absolute deviation from a straight trajectory and movement times. Like pre-movement reaction times, these during-movement measures of movement time and curvature are also sensitive to decision-difficulty, with harder decisions resulting in longer duration movements and greater trajectory curvature (as seen in Figure 1 and [28, 26, 17, 20, 44, 31, 45]).

**Figure 1.**
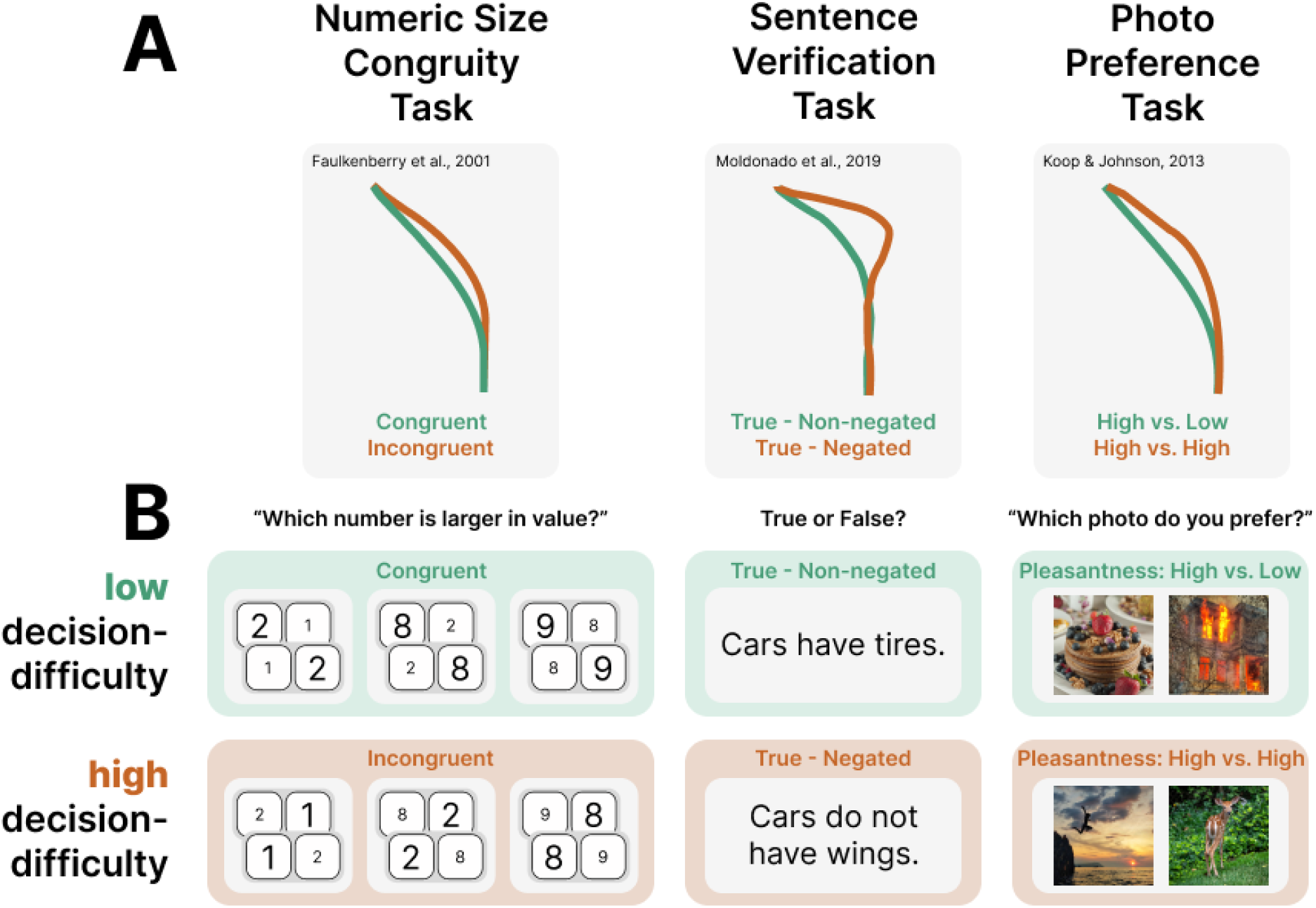
A) From left to right, a recreation of previous mouse trajectory results from the three task we replicate. Shown are average trajectories for the low (green) and high (orange) decision-difficulty categories for the Numeric-Size Congruity task (adapted from [17]), the Sentence Verification task (adapted from [31]’s replication of [13]), and the Photo Preference task (adapted from [28]). B) A representation of trial conditions falling within the low (green shading) and high (orange shading) decision-difficulty categories for each task, with stimuli examples.

Despite reach-decision trajectory-tracking being an important tool for the understanding of decision-making, these approaches remain relatively unused outside of research labs. Recognizing that research deployed online via portable devices could reach a wider and more diverse audience, there has been a recent movement to assess the reliability of cognitive task administration in these environments [2, 39, 37]. This has been fuelled by new tools allowing the development of online tasks (e.g., Labvanced [18], Gorilla [3], jsPsych [30]) that include easy deployment to diverse, crowd-sourced participant pools (e.g MTurk [1], Prolific [35]) and can target a variety of devices [2].

While cognitive tasks measuring accuracy and reaction time have been replicated on tablets [19, 42] and smartphones [4], it is largely unknown if and how motoric measures of decision-difficulty can be measured on these portable devices. To test this question, we developed a reach-decision task using Labvanced [18] to collect continuous cursor position data, and deployed it to over 300 crowd-sourced participants. Critically, each of these participants completed the task on one of three different devices (*>*100 participants per device) varying in size and user-interaction requirements: personal computers (mouse-based interactions), tablets (finger or stylus-based interactions) and smartphones (finger-,thumb- or stylus-based interactions).

To provide evidence that a particular device is tracking decision-difficulty, we chose to replicate three unique reach-decision tasks. Each of these tasks has been shown to sensitively reflect decision-difficulty effects through mouse-tracking (see Figure 1A) and here we tested if those effects were replicable and then extensible to tablets and smartphones. The three tasks were: a Numeric-Size Congruity task [17], a Sentence Verification task [13] and a Photo Preference task [28]. Based on these previous publications, we were able to select trials in each task that reflected high decision-difficulty or low decision-difficulty choices (see Figure 1B). This established a clear benchmark for replication: a particular device was sensitive to decision-difficulty if high decision-difficulty trials displayed significantly greater reaction time, movement time and trajectory curvature scores compared to low decision-difficulty trials [28, 13, 17].

In the Numeric-Size Congruity task, participants were asked to select which of two digits was larger in value, with the paired digits being either congruent in numeric and physical size (low decision-difficulty, e.g., 2 vs. 8) or incongruent in numeric and physical size (high decision-difficulty, e.g., 2 vs. 8). The Sentence Verification task asked participants to verify the truth of statements that could be non-negated (low decision-difficulty, e.g., ‘Cars have tires’) or negated (high decision-difficulty, e.g., ‘Cars do not have wings’). Finally, the Photo Preference task asked participants to select which of two dissimilarly-valenced (low decision-difficulty, e.g., High vs. Low pleasantness) or similarly-valenced (high decision-difficulty, e.g., High vs. High pleasantness) photos they preferred. Together, we ensured these tasks spanned a range of decision domains from objective perceptual judgments (e.g., digit discrimination), to semi-subjective conceptual judgements (e.g., truth value of a statement), and finally subjective preference judgements (e.g., preference for a particular photograph). These tasks also intentionally differed in stimulus characteristics (e.g., numeric, alphabetic, image), stimuli (e.g., numerical digits, written statements, photos), and processing requirements (e.g., perceptual discrimination, conceptual discrimination) allowing our results to be generalizable across remarkably distinct decision domains. Moreover, our experimental design allowed for a thorough exploration of the consistency of, and relationships between, metrics of decision-difficulty at different time points in the decision process (e.g., before and after movement-initiation). Finally, by building on previous mouse-tracking studies we are able to make strong a-priori predictions to provide a definitive test for using widely available touch-devices as a means of vastly extending the reach of decision science.

## 2 Results

### 2.1 Tablets and smartphones measure decision-difficulty as well as computer mouse-tracking during reach-decision tasks

For all three tasks decision-difficulty was quantified as standardized reaction time, movement time and trajectory curvature (MAD) scores (see Methods subsection 5.3.2 - Dependent Measures). A replication of difficulty-driven effects was considered to have occurred should high decision-difficulty trials display significantly greater standardized scores than low decision-difficulty trials [28, 13, 17]. Thus, for each device (computer, tablet, smartphone) a-priori comparisons (t-tests) were made between high and low decision-difficulty trials within each task. A summary of statistics, unstandardized means, and mean differences between standardized scores are reported in Table 1.

**Table 1:** Task-specific unstandardized and z-scored means, and a-priori comparison results. Note. *p *<* .05; **p *<* .005; ***p *<* .0005

| Device | Unstandardized |  |  |  |  |  | Standardized |  |  |  |  |  |
| --- | --- | --- | --- | --- | --- | --- | --- | --- | --- | --- | --- | --- |
|  | *M* |  | *SD* |  |  |  | *Z*<sub>Hard</sub>-*Z*<sub>Easy</sub> |  |  |  |  |  |
|  | *Decision Difficulty* |  | Within |  | Between |  |  |  |  |  |  |  |
|  | Easy | Hard | Easy | Hard | Easy | Hard | *M* | *SE* | df | *t* | *p* | Cohen's *d* |
| Numeric-Size Congruity |  |  |  |  |  |  |  |  |  |  |  |  |
| Reaction Time (ms) |  |  |  |  |  |  |  |  |  |  |  |  |
| Computer | 530.19 | 564.46 | 187.73 | 189.06 | 245.15 | 264.11 | 0.24 | 0.025 | 82 | 9.77 | *** | 1.07 |
| Tablet | 556.12 | 571.19 | 127.02 | 124.66 | 149.23 | 152.21 | 0.16 | 0.026 | 78 | 6.14 | *** | 0.69 |
| Smartphone | 503.24 | 526.08 | 121.35 | 135.78 | 169.16 | 184.01 | 0.18 | 0.034 | 77 | 5.15 | *** | 0.58 |
| Movement Time (ms) |  |  |  |  |  |  |  |  |  |  |  |  |
| Computer | 413.22 | 422.91 | 124.87 | 135.69 | 116.09 | 116.70 | 0.09 | 0.027 | 82 | 3.45 | ** | 0.39 |
| Tablet | 513.31 | 538.46 | 85.38 | 102.68 | 135.83 | 146.86 | 0.27 | 0.029 | 78 | 9.35 | *** | 1.05 |
| Smartphone | 469.64 | 498.98 | 89.62 | 109.12 | 120.13 | 119.29 | 0.31 | 0.029 | 77 | 10.55 | *** | 1.20 |
| Maximum Absolute Deviation (px) |  |  |  |  |  |  |  |  |  |  |  |  |
| Computer | 8.01 | 19.64 | 37.10 | 53.57 | 18.85 | 23.86 | 0.24 | 0.029 | 82 | 8.24 | *** | 0.90 |
| Tablet | 13.04 | 25.75 | 27.70 | 38.18 | 10.55 | 13.42 | 0.38 | 0.030 | 78 | 12.51 | *** | 1.41 |
| Smartphone | 12.90 | 26.92 | 30.72 | 41.87 | 8.93 | 14.10 | 0.37 | 0.027 | 77 | 13.94 | *** | 1.58 |
| Sentence Verification |  |  |  |  |  |  |  |  |  |  |  |  |
| Reaction Time (ms) |  |  |  |  |  |  |  |  |  |  |  |  |
| Computer | 961.78 | 1496.46 | 326.40 | 493.04 | 395.28 | 631.25 | 1.15 | 0.043 | 82 | 26.57 | *** | 2.92 |
| Tablet | 1013.20 | 1403.15 | 312.79 | 489.69 | 344.36 | 596.18 | 0.81 | 0.056 | 78 | 14.43 | *** | 1.62 |
| Smartphone | 1041.42 | 1448.34 | 340.22 | 508.72 | 414.00 | 802.53 | 0.82 | 0.061 | 77 | 13.33 | *** | 1.51 |
| Movement Time (ms) |  |  |  |  |  |  |  |  |  |  |  |  |
| Computer | 462.91 | 606.26 | 174.68 | 274.51 | 164.11 | 257.50 | 0.52 | 0.050 | 82 | 10.50 | *** | 1.15 |
| Tablet | 686.74 | 1056.88 | 215.76 | 413.78 | 241.59 | 631.90 | 0.79 | 0.055 | 78 | 14.26 | *** | 1.61 |
| Smartphone | 627.03 | 995.52 | 199.39 | 413.78 | 210.84 | 491.94 | 0.91 | 0.050 | 77 | 18.40 | *** | 2.08 |
| Maximum Absolute Deviation (px) |  |  |  |  |  |  |  |  |  |  |  |  |
| Computer | 16.09 | 30.09 | 37.90 | 56.15 | 31.13 | 43.45 | 0.25 | 0.048 | 82 | 5.07 | *** | 0.56 |
| Tablet | 16.70 | 35.64 | 27.31 | 36.12 | 27.79 | 42.43 | 0.44 | 0.054 | 78 | 8.13 | *** | 0.91 |
| Samrtphone | 7.06 | 28.12 | 32.39 | 41.64 | 32.02 | 41.78 | 0.48 | 0.059 | 77 | 8.13 | *** | 0.92 |
| Photo Preference |  |  |  |  |  |  |  |  |  |  |  |  |
| Reaction Time (ms) |  |  |  |  |  |  |  |  |  |  |  |  |
| Computer | 1024.93 | 1195.25 | 377.06 | 529.52 | 431.12 | 607.31 | 0.30 | 0.046 | 82 | 6.49 | *** | 0.71 |
| Tablet | 1012.67 | 1048.80 | 389.61 | 379.82 | 454.26 | 576.58 | 0.04 | 0.048 | 78 | 0.80 | n.s. | 0.09 |
| Smartphone | 930.96 | 983.36 | 319.14 | 349.04 | 594.18 | 627.82 | 0.08 | 0.046 | 77 | 1.72 | n.s. | 0.20 |
| Movement Time (ms) |  |  |  |  |  |  |  |  |  |  |  |  |
| Computer | 569.72 | 648.63 | 219.20 | 268.96 | 291.89 | 501.46 | 0.16 | 0.040 | 82 | 3.90 | *** | 0.43 |
| Tablet | 782.13 | 895.75 | 235.46 | 325.69 | 298.36 | 398.20 | 0.23 | 0.047 | 78 | 4.95 | *** | 0.56 |
| Smartphone | 722.06 | 796.08 | 231.83 | 308.75 | 258.66 | 373.29 | 0.16 | 0.044 | 77 | 3.28 | * | 0.37 |
| Maximum Absolute Deviation (px) |  |  |  |  |  |  |  |  |  |  |  |  |
| Computer | 15.43 | 22.41 | 34.72 | 47.05 | 33.65 | 38.87 | 0.15 | 0.045 | 82 | 3.43 | ** | 0.38 |
| Tablet | 20.90 | 30.89 | 33.85 | 36.08 | 21.71 | 20.35 | 0.32 | 0.057 | 78 | 5.87 | *** | 0.64 |
| Smartphone | 24.16 | 31.056 | 32.61 | 38..31 | 25.06 | 21.39 | 0.15 | 0.054 | 77 | 2.78 | * | 0.32 |

For the Numeric-Size Congruity and Sentence Verification tasks, the paired samples t-tests replicated difficulty-driven for all three devices and for all three measures of decision-difficulty (see Table 1 and Figures 2 and 3). The Photo Preference task similarly replicated expected difficulty-driven effects across all measures during computer use, as well as for movement time and trajectory curvature during tablet and smartphone use (see Table 1 and Figure 4). Together, these results suggest that tablets and smartphones are sensitive tools for capturing information-rich reach-decision data across a variety of decision domains. Given the consistency of results for the other two tasks we attribute the divergence between computer and touch-device reaction time results during Photo Preference decisions to task features. Only the Photo Preference task required the judgment of a picture and we believe the fidelity of the picture information is degraded as screen-size is reduced, driving down the sensitivity to difficulty-driven effects on smaller displays. The relative increase in sensitivity to decision-difficulty for Computer reaction times is consistent with the Device differences described in the next Results subsection.

**Figure 2.**
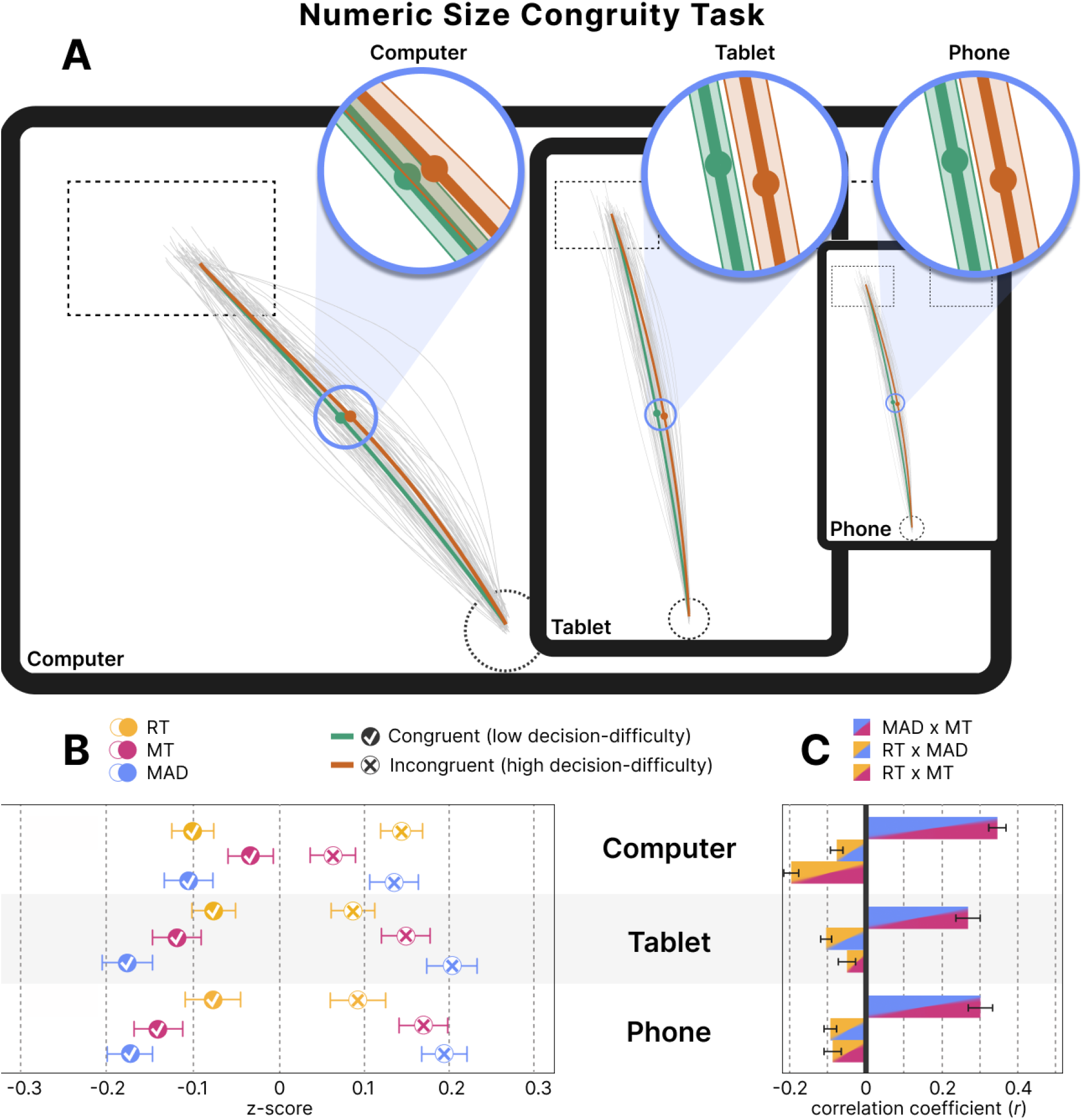
Numeric-Size Congruity task results. A) From left to right, trajectory results for computer, tablet and smartphone (phone) devices within screen size boundaries shown to scale of a representative physical device size. Light gray lines are each participants’ average trajectory across all trials in this comparison. Mean trajectories across participants are shown for low (green line, Congruent trials) and high (orange line, Incongruent trials) decision-difficulty trials with the average location of maximum absolute deviation (MAD) shown with a filled circle. Insets zoom-in on the average point of MAD. Rightward reaches were mirrored to end left, and all reaches were space-normalized and standardized. Errors shown in the insets are the average of within-subjects standard error. For full trajectory visualization details, see Supplementary Note 1. B) From top to bottom, average of participant mean z-scored reaction times (yellow), movement times (pink), and maximum absolute deviation (blue) for computer, tablet and smartphone use. Error bars represent the averaged standard error of the difference between high and low difficulty means. C) Pearson’s correlations (*r*) between measures of decision-difficulty for (from top to bottom) computer, tablet and smartphone use calculated from each participant and shown as an average. Error bars represent the standard error of the estimated marginal mean.

**Figure 3.**
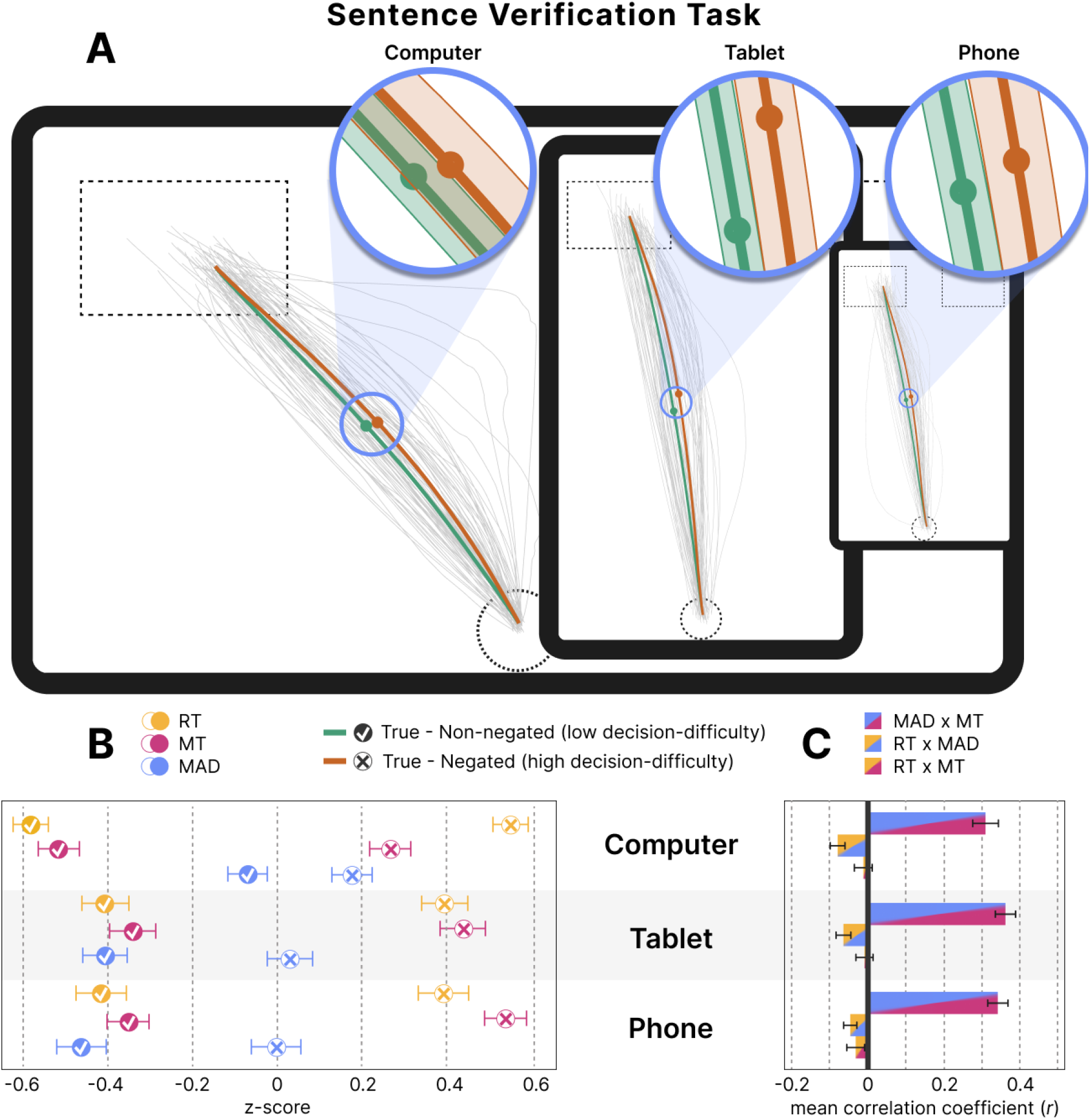
Sentence Verification task results. A) From left to right, trajectory results for computer, tablet and smartphone (phone) devices within screen size boundaries shown to scale of a representative physical device size. Light gray lines are each participants’ average trajectory across all trials in this comparison. Mean trajectories across participants are shown for low (green line, True Non-negated trials) and high (orange line, True Negated trials) decision-difficulty trials with the average location of maximum absolute deviation (MAD) shown with a filled circle. Insets zoom-in on the average point of MAD. Rightward reaches were mirrored to end left, and all reaches were space-normalized and standardized. Errors shown in the insets are the average of within-subjects standard error. For full trajectory visualization details, see Supplementary Note 1. B) From top to bottom, average of participant mean z-scored reaction times (yellow), movement times (pink), and maximum absolute deviation (blue) for computer, tablet and smartphone use. Error bars represent the averaged standard error of the difference between high and low difficulty means. C) Pearson’s correlations (*r*) between measures of decision-difficulty for (from top to bottom) computer, tablet and smartphone use calculated from each participant and shown as an average. Error bars represent the standard error of the estimated marginal mean.

**Figure 4.**
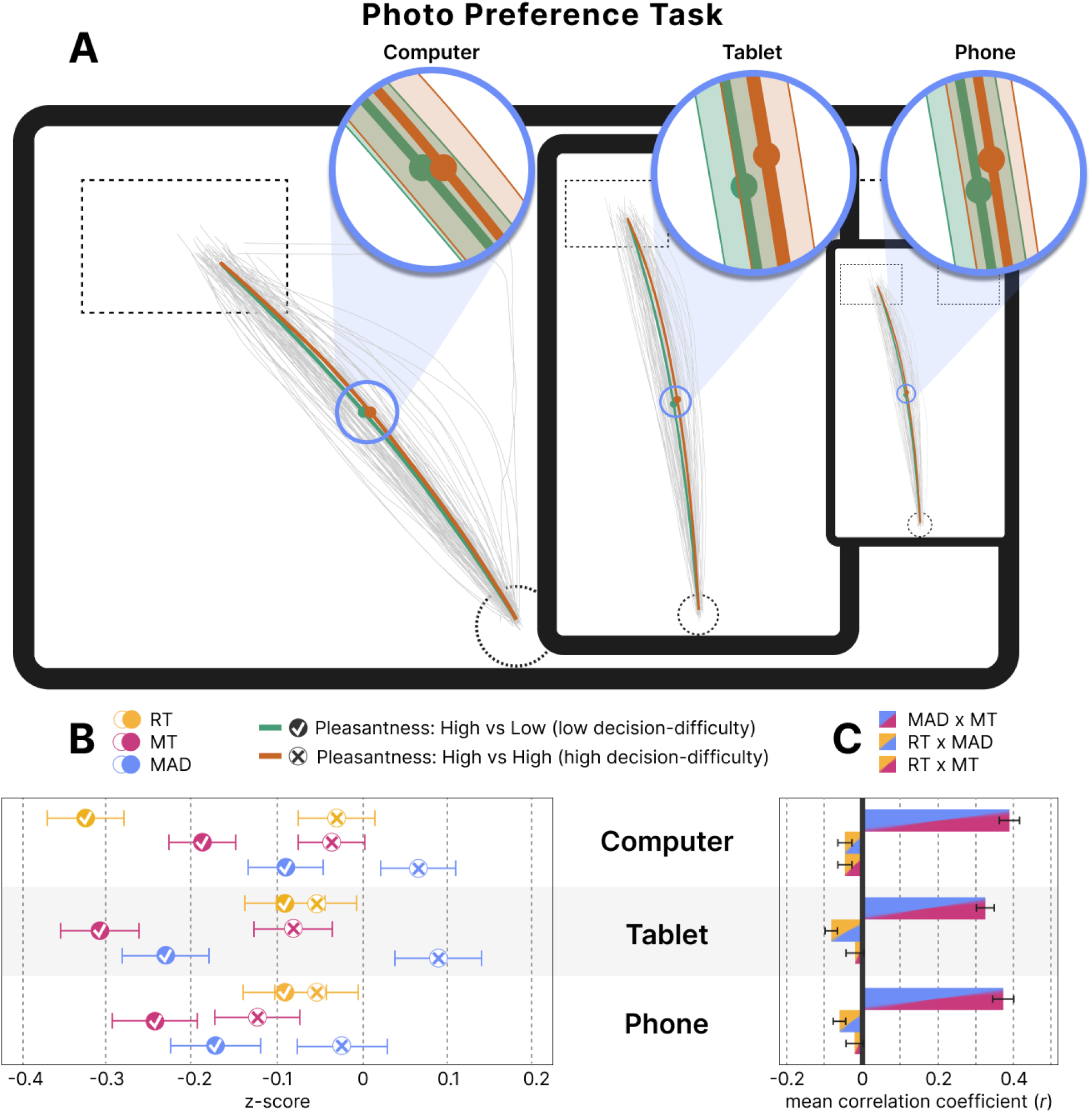
Photo Preference task results. A) From left to right, trajectory results for computer, tablet and smartphone (phone) devices within screen size boundaries shown to scale of a representative physical device size. Light gray lines are each participants’ average trajectory across all trials in this comparison. Mean trajectories across participants are shown for low (green line, High vs. Low pleasantness trials) and high (orange line, High vs. High pleasantness trials) decision-difficulty trials with the average location of maximum absolute deviation (MAD) shown with a filled circle. Insets zoom-in on the average point of MAD. Rightward reaches were mirrored to end left, and all reaches were space-normalized and standardized. Errors shown in the insets are the average of within-subjects standard error. For full trajectory visualization details, see Supplementary Note 1. B) From top to bottom, average of participant mean z-scored reaction times (yellow), movement times (pink), and maximum absolute deviation (blue) for computer, tablet and smartphone use. Error bars represent the averaged standard error of the difference between high and low difficulty means. C) Pearson’s correlations (*r*) between measures of decision-difficulty for (from top to bottom) computer, tablet and smartphone use calculated from each participant and shown as an average. Error bars represent the standard error of the estimated marginal mean.

### 2.2 Mouse-tracking is more sensitive to decision-difficulty before movement while touch-device interactions are more sensitive during movement

Having established that all three devices tested capture decision-difficulty, our second analyses tested *how* the measurement of decision-difficulty changed across devices. Mean standardized reaction times, movement times and trajectory curvature scores for each task were separately submitted to a mixed-model ANOVA where we focused on main effects or interactions involving the between-subjects factor of Device factor and explored any (simple) main effects with pairwise comparisons between levels of Device (for results from this analysis outside this specific scope, including those that fully support the a-priori decision-difficulty effects described above, see Supplementary Materials 1). These tests revealed that the sensitivity of the specific metrics of decision-difficulty differed between touch-device and computer interactions. Specifically, computers showed increased sensitivity to decision-difficulty pre-movement (i.e., reaction time) while tablets and smartphones showed increased sensitivity during movement (i.e., movement time and trajectory curvature).

#### 2.2.1 Measure sensitivity pre-movement

Within the Numeric-Size Congruity task, a 2 (Congruity) x 3 (Number Pairs) x 2 (Number Presentation Side) x 3 (Device) mixed-model ANOVA assessing standardized reaction times revealed both a main effect of Device (*F* (2,237) = 12.69, *p* = 5.81e-6, *η*^2^ = 3.16e-4) and an interaction between Number Pair and Device (*F* (4,237) = 14.23, *p* = 3.37e-10, *η*^2^ = .022). A significant main effect of Device was seen for both 1v2 (*F* (2,237) = 17.79, *p* = 6.31e-8) and 8v9 Number Pairings (*F* (2,237) = 19.77, *p* = 1.15e-8). The 8v9 effect, which is the hardest number-pair to decide between because it has both the smallest numeric difference and the smallest relative difference (see Supplementary Discussion 2), is driven by Computer having the longest reaction times compared to the touch-devices (*Mean*_*Computer−Smartphone*_ = 0.18, *t* = 5.74, *p* = 6.01e-7, *d* = 0.43; *Mean*_*Computer−Tablet*_ = 0.20, *t* = 6.78, *p* = 1.30e-9, *d* = 0.50). Meanwhile, the 1v2 effect, which is much easier because of the larger relative difference and presence of small numbers, is driven by Computer having the shortest reaction times (*M* _*Computer−Smartphone*_ = -0.13, *t* = 4.26, *p* = 8.77e-4, *d* = 0.32 and *M* _*Computer−Tablet*_ = -0.16, *t* = 5.26, *p* = 7.74e-6, *d* = 0.39). Thus, for reaction time, Computers show greater differentiation between hard and easy trials.

A similar pattern emerged in the Sentence Verification task. A 2 (Truth Value) x 2 (Polarity) x 3 (Device) mixed-model ANOVA revealed a three way interaction Truth x Negation x Device (*F* (2,237) = 8.21, *p* = 3.57e-4, *η*^2^ = .005) within reaction time. Based on where we predicted decision-difficulty to differ (see Figure 1) our follow-up tests looked at Negation x Device for True and False statements. We found a significant interaction only for True statements (*F* (2,237) = 13.32, *p* = 3.30e-6, *η*^2^ = .022). Breaking this down, Device was significant for both True-Negated statements (*F* (2,238) = 8.22, *p* = 3.55e-4) and True-Non-negated statements (*F* (2,238) = 14.27, *p* = 1.40e-6), but in importantly different ways. For the more difficult True-Negated statements, Computer reaction times were the longest (*M* _*Computer−Tablet*_ = 0.16, *t* = 3.76, *p* = .003, *d* = 0.59; *M* _*Computer−Smartphone*_ = 0.16, *t* = 3.82, *p* = .002, *d* = 0.60), but, for the easier True-Non-Negated statements, Computer reaction times were the shortest (*M* _*Computer−Tablet*_ = -0.18, *t* = 4.25, *p* = 4.19e-4, *d* = 0.67; *M* _*Computer−Smartphone*_ = -0.17, *t* = 4.11, *p* = 7.53e-4, *d* = 0.65). These results confirm that computers show greater differentiation across levels of decision-difficulty.

#### 2.2.1 Measure sensitivity during-movement

An opposite pattern of results can be found when analyzing standardized movement time. Using the same ANOVA model described above, for Numeric-Size Congruity we found an interaction between Congruity and Device (*F* (2,237) = 16.51, *p* = 1.93e-7, *η*^2^ = .009). Follow-ups showed Device was significant for both Congruent (*F* (2,237) = 18.15, *p* = 4.63e-8) and Incongruent trials (*F* (2,237) = 14.22, *p* = 1.47e-6). Here, Computer showed *increased* movement times for Congruent trials (*M* _*Computer−Smartphone*_ = 0.11, *t* = 5.38, *p* = 2.61e-6, *d* = 0.26; *M* _*Computer−Tablet*_ = 0.088, *t* = 4.34, *p* = 3.06e-4, *d* = 0.21) but *decreased* movement times for Incongruent trials (*M* _*Computer−Smartphone*_ = -0.11, t = 5.20, *p* = 6.08e-6, *d* = 0.21; *M* _*Computer−Tablet*_ = -0.087, *t* = 4.30, *p* = 3.67e-4, *d* = 0.21), resulting in less divergence in movement times between the two difficulty levels compared to touch-devices. In complete opposition to the pattern observed for reaction times, these results suggest Computer movement times are significantly less sensitive to decision-difficulty compared to Tablet and Smartphone movement times.

Again Sentence Verification movement time results confirm this finding. Here the same task-specific mixed-model ANOVA described previously revealed a Negation by Device interaction (*F* (2,237) = 19.59, *p* = 1.34e-8, *η*^2^ = .027). Follow-ups revealed a main effect of Device both when statements were Non-negated (*F* (2,237) = 21.43, *p* = 2.78e-9) and Negated (*F* (2,237) = 16.82, *p* = 1.48e-7). Pairwise comparisons showed Computer having longer movement times compared to Tablets and Smartphones when statements were Non-negated (*M* _*Computer−Smartphone*_ = 0.15, *t* = 5.76, *p* = 3.53e-7, *d* = 0.57; *M* _*Computer−Tablet*_ = 0.12, *t* = 4.54, *p* = 1.33e-4, *d* = 0.44) and shorter movement times when statements were Negated (*M* _*Computer−Smartphone*_ = -0.15, *t* = 5.96, *p* = 1.20e-7, *d* = 0.59; *M* _*Computer−Tablet*_ = -0.11, *t* = 4.29, *p* = 3.85e-4, *d* = 0.42). This again results in less sensitivity in movement time between levels of Negation for the Computer condition compared to touch-devices.

The during-movement sensitivity observed for touch-devices also extended to trajectory curvature, but was impacted by the biomechanical properties of using a hand to act directly on a screen. Specifically, both tablet and smartphone results displayed a side of space biases where rightward reaches show more trajectory curvature compared to leftward reaches, matching what is observed in real reaching experiments [21]. Within Numeric-Size Congruity, this effect is evident in the trajectory curvature results as a Number Pair Presentation Side x Device interaction (*F* (2,237) = 16.90, *p* = 1.38e-7, *η*^2^ = .049) where both Left and Right reaches showed main effects of Device (Left: *F* (2,237) = 17.07, *p* = 1.19e-7; Right: (*F* (2,237) = 16.55, *p* = 1.86e-7), but in opposite directions. For Left reaches, Tablets and Smartphones show significantly less curvature than Computer trajectories (*M* _*Computer−Tablet*_ = 0.27, *t* = 4.70, *p* = 6.47e-5, *d* = 0.52; *M* _*Computer−Smartphone*_ = 0.30, *t* = 5.34, *p* = 3.30e-6, *d* = 0.59) while for Right reaches, Tablets and Smartphones show significantly more curvature than Computer trajectories (*M* _*Computer−Tablet*_ = -0.26, *t* = 4.66, *p* = 7.96e-5, *d* = 0.51; *M* _*Computer−Smartphone*_ = -0.29, *t* = 5.20, *p* = 6.57e-6, *d* = 0.57). Appreciating that Sentence Verification choice stimuli were locked to a side of space, the Sentence Verification trajectory curvature results bolster these directional effect findings, revealing a Truth x Device interaction (*F* (2,237) = 15.16, *p* = 6.39e-7, *η*^2^ = .074). Here we also see main effects of Device for both Left/True (*F* (2,237) = 13.96, *p* = 1.86e-6) and Right/False reaches (*F* (2,237) = 16.23, *p* = 22.47e-7) but in opposite directions. For Left/True reaches, Tablets and Smartphones show significantly less curvature than Computer trajectories (*M* _*Computer−Tablet*_ = 0.25, *t* = 4.28, *p* = 4.06e-4, *d* = 0.59; *M* _*Computer−Smartphone*_ = 0.30, *t* = 5.10, *p* = 1.03e-5, *d* = 0.70) while for Right/False reaches, Tablets and Smartphones show significantly more curvature than Computer trajectories (*M* _*Computer−Tablet*_ = -0.24, *t* = 4.18, *p* = 6.12e-4, *d* = 0.58; *M* _*Computer−Smartphone*_ = -0.30, *t* = 5.09, *p* = 1.06e-5, *d* = 0.70). Again, this suggests that a right hand bias is more prominent for real touch interactions compared to mouse cursor movements (see Supplemental Discussion 3 for confirmatory evidence from the analysis of Movement Time).

Finally, the trajectory results from the Photo Preference task provide another example of how touch and mouse interactions differ. A 3 (Valence Pairing) x 3 (Device) mixed-model ANOVA revealed a main effect of Device (*F* (2,237) = 9.32, *p* = 1.27e-4, *η*^2^ = .022) with standardized trajectory values for Computer responses (*M* = -0.0263, *SD* = 0.267) found to be different than Tablet (*M* = -0.116, *SD* = 0.322; *t* = 3.50, *p* = .001, *d* = 0.31) and Smartphone responses (*M* = -0.132, *SD* = 0.036; *t* = 3.91, *p* = 3.64e-4, *d* = 0.35), and no significant difference between the two touch-devices. This Device effect did not significantly interact with decision-difficulty, indicating that this is a difference in the shape of the produced trajectories based on input - an idea which aligns with our interpretation that reaches produced as a result of direct interaction are different than those mediated by a mouse (see section 3 - Discussion). Overall, the differences in trajectory shape and presence of a right-hand bias in the Tablet and Smartphone results in contrast to Computer results point to a similarity between touch-device responses and real-world reaching when making choice selections. Further, these results highlight the increased sensitivity of post-movement measures during touch-device use.

### 2.3 Pre- and post-movement measures are flexible, non-redundant carriers of decision information

Here, we assess the relationship between our decision-difficulty measures to demonstrate that pre- and during-movement measures carry unique decision information. To do so, we obtained a within-participant correlation coefficient (*r*) for each combination of measures (Correlation-Type: *r* _*MAD,MT*_ vs. *r* _*MAD,RT*_ vs. *r* _*MT,RT*_) within each task and device. These participant average correlation coefficients were then compared using a (3) Correlation Type x (3) Task x (3) Device mixed-model ANOVA. Where correlations between measures are positive, it would indicate that they carry redundant information. However, any inverse relationship would demonstrate a push and pull between measures showing that on any given trial, a best estimate of decision-difficulty should include both pre- and during-movement measures. The results of the ANOVA revealed a main effect of Task (*F* (2,237) = 22.06, *p* = 1.13e-9, *η*^2^ = .009), a very strong main effect of Correlation-Type (*F* (2,237) = 601.10, *p* = 1.10e-92, *η*^2^ = .45) and an interaction between Correlation-Type and Task (*F* (4,237) = 5.54, *p* = 6.47e-7, *η*^2^ = .004). To follow up, we examined each Task separately and found a strong Correlation-Type effect in all three (SC: *F* (2,239) = 302.94, *p* = 2.85e-69, *η*^2^ = .56; SV: *F* (2,239) = 242.55, *p* = 6.05e-53, *η*^2^ = 0.50; PP: *F* (2,239) = 358.29, *p* = 6.13e-76, *η*^2^ = .60). Mean *r* values revealed trajectory curvature and movement time (*r* _*MAD,MT*_) to be moderately positively correlated (SC: *M* _*r*_ = 0.30, *SD* = 0.24; SV: *M* _*r*_ = 0.33, *SD* = 0.26; PP: *M* _*r*_ = 0.36, *SD* = 0.23) which intuitively makes sense - traveling a longer distance (MAD) usually takes a longer time (MT). In contrast, in each task, reaction time was found to be weakly inversely correlated with both other measures (SC: *M* _*r*_ = -0.092, *SD* = 0.14 and *M* _*r*_ = -0.11, *SD* = 0.20 for *r* _*MAD,RT*_ and *r* _*MT,RT*_ correlations, respectively; SV: *M* _*r*_ = -0.065, *SD* = 0.17 and *M* _*r*_ = 0.006, *SD* = 0.20 for *r* _*MAD,RT*_ and *r* _*MT,RT*_ correlations, respectively; PP: *M* _*r*_ = -0.065, *SD* = 0.15 and *M* _*r*_ = -0.041, *SD* = 0.19 for *r* _*MAD,RT*_ and *r* _*MT,RT*_ correlations, respectively). This pattern meant that the Correlation-Type comparisons always showed differences between during-movement correlations (*r* _*MAD,MT*_, stronger and positive) and the pre-to during-movement correlations (*r* _*MAD,RT*_ and *r* _*MT,RT*_, weaker and negative). By task, the results of these pairwise comparisons were, for *r* _*MAD,MT*_ vs. *r* _*MAD,RT*_ : SC: *p* = 3.5e-68, *d* = 2.00; SV: *p* = 1.06e-67, *d* = 1.86; PP: *p* = 8.23e-83, *d* = 2.19, and for *r* _*MAD,MT*_ vs. *r* _*MT,RT*_ : SC: *p* = 2.18e-73, *d* = 2.11; SV: *p* = 2.15e-50, *d* = 1.53; PP: *p* = 2.04e-76, *d* = 2.07. The only slight difference across tasks we observed was that *r* _*MT,RT*_ in the Sentence Verification task was close to zero, rather than weakly negative, and as such, there was a pairwise difference between *r* _*MT,RT*_ and *r* _*MAD,RT*_ (*p* = 7.70e-04, *d* = -0.33).

Taken together, this analysis reveals that pre- and during-movement measures display an intricate relationship independent of their role in indexing task-specific decision-difficulty. That is, while across all tasks and devices, reaction time, movement time and curvature increase with decision-difficulty (see Results subsection 2.1) on a trial-by-trial basis these measures adapt to the demands of the task and pre- and during-movement measures function as non-redundant carriers of decision information. Specifically, it appears that on trials where participants react more quickly (shorter RTs) there is a slight increase in movement time and curvature (see section 3 - Discussion for further interpretation). It is also notable that there were no significant Device differences and limited differences due to Task. This highlights the remarkable stability both of this interplay between measures and for reach-decisions to track decision-difficulty across a variety of interface types.

## 3 Discussion

We investigated whether measuring reach decision-difficulty could be extended beyond computer use to tablets and smartphones through the deployment of a three-task online experiment across the three devices. Each task replicated a prior mouse-tracking study used to observe decision processes (Numeric-Size Congruity task [17], Sentence Verification task [13, 31], Photo Preference task [28]), allowing us to make strong predictions about which trials in each task would have high versus low decision-difficulty (see Figure 1).

Task-specific results replicated previous mouse-tracked outcomes, with high difficulty decisions displaying greater reaction times, movement times and trajectory curvature compared to low difficulty decisions. Most excitingly, all of these effects were replicated across all devices. Thus, this study demonstrates the robustness of dynamic measures of decision-making and offers validation for the use of small, portable devices to collect this movement information. For the Numeric-Size Congruity task [17], replication manifested as increased reaction time, movement time and trajectory curvature for incongruent trials compared to congruent trials (see Figure 2). For the Sentence Verification task [13, 31], the same metrics were increased on true-negated statements compared to true-non-negated statements (see Figure 3). Finally, for the Photo Preference task [28], movement time and trajectory curvature were increased for decisions requiring judgements between photos similar in pleasantness compared to decisions requiring judgements between photos dissimilar in pleasantness.

However, these a-priori comparisons also suggested that not all tasks might be suitable for deployment on smaller devices. Results from the Photo Preference task show that tablets and smartphones have a reduced sensitivity to decision-difficulty effects, especially for reaction time (see Table 1). We believe that this is a reflection of stimuli salience as screen size is reduced. While the other two tasks presented decision information as text, the Photo Preference task required participants to distinguish between two detailed photos, which likely degraded in stimulus information as the stimulus size decreased. Therefore, our key message is that all devices are able to track decision-difficulty but device differences exist and are important to understand. Our second cluster of results then specifically interrogated device differences. The results were clear: computer responses were consistently different from tablet and smartphone responses. Computer responses showed an increased sensitivity to decision-difficulty within pre-movement measures (reaction time) while touch-device responses revealed greater sensitivity during movement (movement time and trajectory curvature). We speculate this might be due to the different user-interaction requirements of touch-devices that enforce different ‘reach’ biomechanics compared to computer-mouse interactions. This is supported by the right-hand bias effects observed when swiping a finger/thumb or sliding a stylus but not when moving a mouse. This right-hand bias, also evident in real reaching [7, 21], is thought to arise from preferential processing of stimuli presented on the right of a display you are interacting with, resulting in less trajectory curvature and faster movement times during rightward reaches.

Why might smartphones and tablets show effects similar to a real reach movement? First, real-world movements made to enact mouse cursor changes on a screen are physically very small. While the cursor traverses a large on-screen distance, the hand moving the mouse travels a smaller distance in less time than even a finger on a smartphone (see non-standardized means in Table 1). These movements across less space and time produce more ballistic responses [24, 23]. As time and space during movement are at a premium with little of either available to express in indecision, this requires more of a decision to be resolved prior to movement initiation. [25, 52, 50]. The repercussions of front-loading the decision due to physical movement constraints align with results demonstrating that the demands of a motor task can directly influence cognitive processing (e.g., cognitive tuning [46, 10, 6, 33]). Here, it means that decision-differences arising in a computer task need to be more resolved prior to movement, leading to more sensitivity to difficulty being expressed by reaction time. More broadly, these results support the idea that the brain is optimized to take advantage of the affordances of the world it navigates, when more time and space are available because a physical movement is longer, the final commitment to a particular choice can be withheld well into movement execution [50].

A second explanation for the difference between pre- and during-movement sensitivity across computers compared to tablets and smartphones is the directness of the interaction. When moving a mouse to control a cursor to select a choice-option the action is physically dissociated from the target we are choosing - the hand is on the table rather than the screen. But, when we move our finger to touch a choice-option on a tablet or smartphone our action is directed toward the actual thing we are selecting. From the perspective of a brain controlling movement this is likely a profoundly different problem. For example, physically interacting with an object increases its appeal [51] and moving an object toward your own body can improve your ability to remember it [48]. These phenomena are likely related to the coordinate remapping required when moving a mouse in one plane to control a cursor in a different plane. This dramatically differs from the more direct planning available to the brain when mapping a touch screen target into the action space of the hand and arm [12, 53, 43, 49]. We would argue that it is this directness of interaction and movements that traverse longer distances over more time that explain why touch-devices show increased sensitivity in measures recorded during movement.

This dynamic interplay between pre- and during-movement measures was the subject of our third category of results. Despite all three measures increasing as decision-difficulty increased, our correlational analyses revealed an inverse relationship between reaction time and during-movement measures (also seen in some previous real-reaching tasks [16]). This discrepancy between overall task-related effects and trial-by-trial effects on the measures is compatible with an evidence accumulation framework of decision-making. Within this framework, evidence is noisily accumulated over time until a decision threshold has been reached [45, 50], signalling the onset of a movement. More difficult decisions require more evidence to be accumulated before support for one option reaches this threshold. This takes more time (i.e., longer reaction time), and unresolved competition impacts movements during choice selection (i.e., longer and less straight movements [47, 45, 50]), explaining the overall effects of decision-difficulty we report. However, when decision-difficulty is constant, there is still natural variation in reaction times. If decision processing requirements remain the same, but reaction time is reduced, there is more unresolved competition at movement onset. This necessarily shifts decision processes into the movement. As a result, on a trial-by-trial basis shorter reaction times will map to longer movement times and trajectories with more curvature - exactly the inverse relationship we report. Evidence accumulation thus accounts for both the a-priori main effects of decision-difficulty we report and the measure correlations we observe. Harder decisions result in increased reaction times, movement times and trajectory curvature because evidence accumulates more slowly in these cases. For any given decision where a set amount of evidence is required, however, there is a trade-off between pre-movement and during-movement decision resolution - abbreviating one elongates the other.

## 4 Conclusion

Across computers, tablets and smartphones, measured by reaction time, movement time and trajectory curvature, and capturing how these measures are dynamically related, reach-decision tasks provide a detailed read-out of decision-making. Given the ubiquitous use of touch-devices and websites, our validation of these metrics - across three diverse tasks and in a remote cohort of 240 participants - prove they are accessible outside the lab and impartial to the device used. The remarkable consistency of our results offers exciting new ways to apply these findings to research and industry, providing detailed knowledge of decision dynamics to domains such as corporate talent assessment and implicit bias measurement. Our results also offer the potential to optimize the collection of decision information, indicating that there are features of a decision and a device that make a certain combination the most sensitive for a particular task. Decisions and the movements we make to enact them literally shape our daily lives. By vastly expanding the accessibility of decision measures to include anyone with a touch-device we therefore hope to open new doors to the insights derived from this rich information.

To build on the incredible opportunity of remote data collection used to investigate the detailed dynamics of decision processes, we also conducted a second companion study (Bertrand et al., 2023). In this subsequent study we replicate the current study but integrate webcam eye-tracking, a technique which is sensitive to pre-movement decision processes. Together, this two-study series allows us a detailed description of the entirety of a decision - from the gaze deployed to gather information upon stimuli onset through the mouse-tracked movements produced to enact a final choice.

## 5 Methods

### 5.1 Participants

All experimental procedures were approved by the University of Alberta Research Ethics Office. 305 naive Amazon Mechanical Turk (www.mturk.com) participants took part in the study using either a computer, tablet or smartphone for a payment of $7 USD. Participation was restricted on Mechanical Turk to Canada- or U.S-based participants between 18 and 35 years of age who had an approval rating above 95% on 100 or more study completions. Participants self-reported age, gender, handedness, visual acuity, English language proficiency, habitual activities requiring hand-eye coordination, chosen device specifications and typical use of their chosen device for participation (see Supplementary Tables 1-3 for a complete demographic and device use summary). Participants were excluded from analysis based on insufficient (*<* 50%) good trials within any of the experimental tasks or in any of the unique task conditions (see sub-subsection 5.3.3 - Data Cleaning).

#### 5.1.1 Computer

101 participants completed the study using a personal computer. Of those, nine were excluded from analysis for not meeting device interaction requirements (i.e., did not use a wired or wireless mouse). A further nine computer users were excluded (see sub-subsection 5.3.3 - Data Cleaning), resulting in data from 83 computer users being analyzed (25 female, 56 male, and 2 who preferred not to say; *M* _*age*_ = 33.75, *SD* _*age*_ = 9.35).

#### 5.1.2 Tablet

101 participants completed the study using a tablet. Four were excluded from analysis for not meeting device interaction requirements (i.e., did not use finger-, thumb-or stylus-based interactions). A further nineteen tablet users were excluded (see sub-subsection 5.3.3 - Data Cleaning), leaving data from 79 tablet users to be analyzed (27 female, 51 male, and 1 nonbinary; *M* _*age*_ = 33.41, *SD* _*age*_ = 6.25).

#### 5.1.3 Smartphone

103 participants completed the study using a smartphone. Of those, twenty-five were excluded (see sub-subsection 5.3.3 - Data Cleaning), leaving 78 smartphone users for analysis (26 female, 52 male, and 1 who preferred not to say; *M* _*age*_ = 33.73, *SD* _*age*_ = 6.72).

### 5.2 Procedure and apparatus

The study was implemented using Labvanced [18], a graphical task builder offering built-in mouse- and finger-tracking, and temporal response recording compatible with computer, tablet and smartphone use for online study implementation. The study was distributed via Amazon Mechanical Turk, and devices used for study completion were uncontrolled except for requiring use of a separate mouse (wired or wireless) during computer use, or an Android operating system and touch-screen device interaction (via finger, thumb or stylus) during tablet or smartphone use (see Supplementary Tables 2-3 for selected device and interaction details).

Participants completed three reach-decision tasks requiring them to choose one of two stimuli presented at the top left and top right corners of their device screen based on a question or statement appearing at the center of the testing interface (see Figure 5). The reach-decision tasks (see Figure 1) presented Numeric-Size Congruity (adapted from [17]), Sentence Verification (adapted from [13, 31]) and Photo Preference (adapted from [28]) paradigms, each consisting of 84 trials and taking approximately 15 minutes to complete.

**Figure 5.**
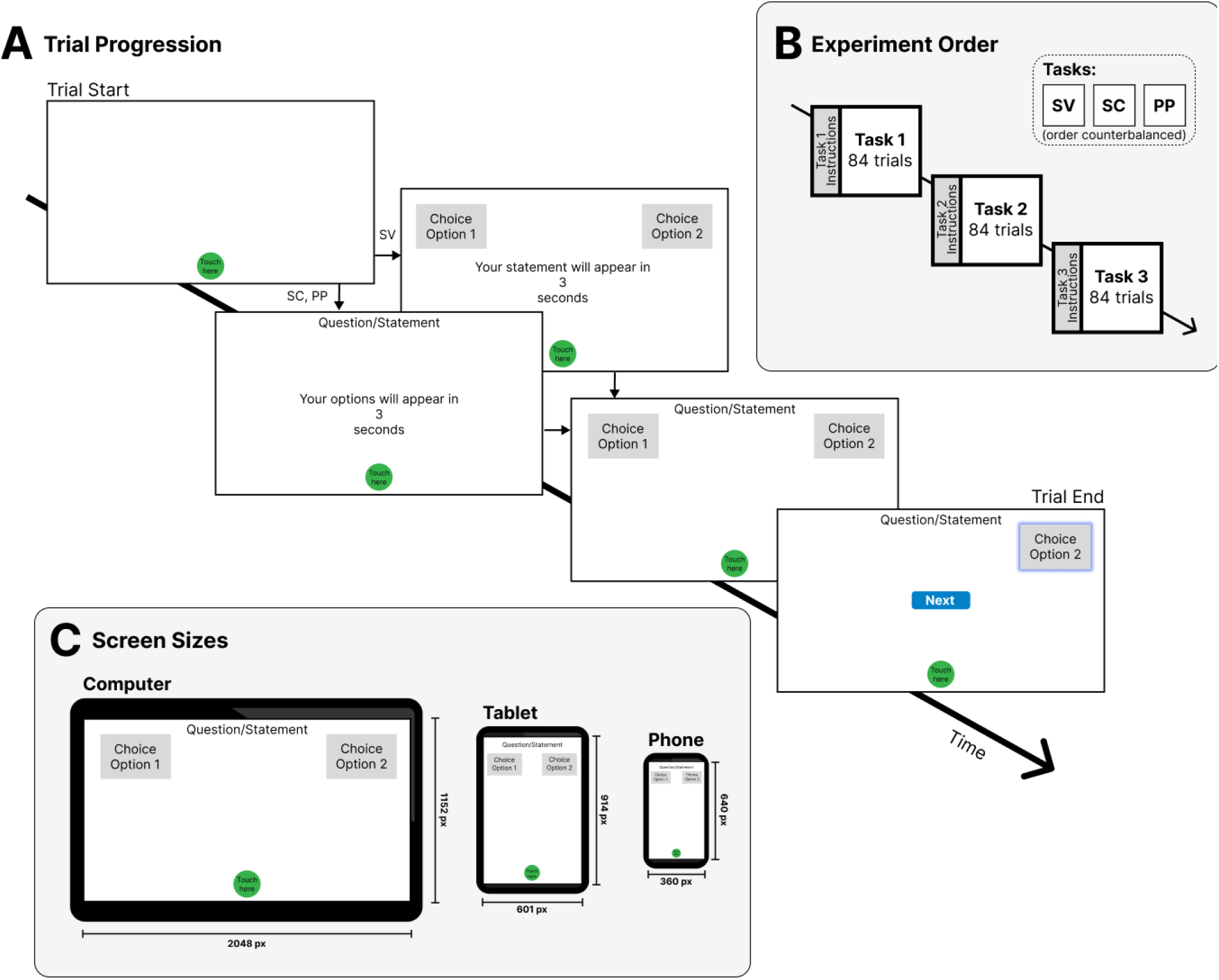
A) Overview of study design. Each participant completed a Numeric-Size Congruity task (SC), a Sentence Verification task (SV) and a Photo Preference Task (PP), with task order counterbalanced between participants. Task-specific instructions were presented prior to each task. B) All three tasks presented a classic reach-decision paradigm requiring participants to choose one of two stimuli presented at the top left and top right of their device screen. For SC and PP tasks, countdown onset was accompanied by a question specific to the task type appearing centered at the top of the display. The SV task presented the two choice options coincident with countdown onset and presented a statement (rather than a question) upon countdown completion. C) A comparison of interface arrangements between devices. Shown are representative examples of a computer, tablet and smartphone (phone) testing interface. All values are reported in pixels. Specific sizes of device screens and interface components observed by participants were dependent on the size of the device used, but screen to interface component proportions remained constant within each device category.

Each trial first presented a green circular start button labeled “Touch here” at the bottom center of the screen, requiring participants to navigate their mouse cursor to (Computer) or place their finger, thumb, or stylus on (Tablet and Smartphone) the button to start the trial. Touching the start button triggered a three second countdown, centered on the display screen (Figure 5B). Removing the mouse cursor, digit or stylus from the start button or the surface of the screen paused the countdown until touch-contact within the start button had been re-established. For the Numeric-Size Congruity and Photo Preference tasks, countdown onset was accompanied by a task-specific question appearing centered at the top of the display (Figure 5B). Upon countdown completion, two choice boxes appeared at the upper-left and upper-right of the screen, each presenting trial-specific choice options. For the Sentence Verification task, the two choice options appeared coincident with countdown onset and presented a statement centered at the top of the screen upon countdown completion (Figure 5B). Participants were free to select either choice option immediately upon countdown completion. For Computers, choice selection required participants to move their mouse cursor inside the choice-box. For Tablets and Smartphones, participants were required to slide their finger, thumb, or stylus across the screen to touch their selected choice-box, keeping contact with the screen at all times. If touchscreen contact was lifted, that trial was removed from analysis and an error message would appear on the screen, reading “Your finger was lifted from the screen as you moved, and we were unable to track the movement. Please touch your option now and remember in the future to keep your finger on the screen.” When selected, a choice-box was highlighted with a blue border, the other option and start button disappeared, and a “Next” button appeared centered on the screen. Participants were then free to click or press on the “Next” button to continue to the next trial, allowing them to self-pace the experiment. Trials were randomized within each task and the order of task presentation was counterbalanced across participants. Participants were instructed to complete the study in its entirety in a single session and were provided with detailed instructions outlining each task before it started. Participants were encouraged to take short breaks between tasks but had a maximum time limit of ninety minutes to complete the study. Labvanced automatically scales the dimensions of the testing interface and its stimuli components to the screen size and resolution of the device in use, presenting a landscape (800 × 450 pixel, Labvanced coordinates) orientation for computer-based participation and a portrait (470 × 800 pixel, Labvanced coordinates) orientation for touch-device based participation. Stimuli-screen proportions remained consistent independent of device screen size (see Figure 5B for device-specific design details).

#### 5.2.1 Numeric-Size Congruity

The Numeric-Size Congruity task in the current study was adapted from Faulkenberry, Cruise, Lavro and Shaki’s experiment [17] examining the dynamics of the size congruity effect. For each Numeric-Size Congruity trial, the question “Which number is larger in value?” appeared coincident with the onset of the countdown timer, centered at the top of the screen (Figure 5). Following countdown termination two numbers were displayed simultaneously, one in each of the upper-left and upper-right choice boxes, and participants could move to select their preferred choice. Stimuli consisted of the Arabic numerals 1, 2, 8 and 9 displayed in Arial font and presented in pairs of different physical size with a 2:1 font size ratio. From these, six choice-pairs were generated: 1v2, 2v8 and 8v9, with each pair either congruent in physical and numeric size (the numerically larger numeral appearing physically larger than its paired counterpart, e.g., 2v8), or incongruent in physical and numeric size (the numerically larger numeral appearing physically smaller than its paired counterpart, e.g., 2v8; see Figure 1). Within each condition, the numerically larger number was presented equally often on the left and the right, counterbalancing side of space effects. This created twelve conditions, each presented 7 times for a total of 84 trials.

#### 5.2.1 Sentence Verification

Adapted from Maldonado, Dunbar and Chemla’s replication [31] of Dale and Duran’s linguistic negation experiment [13], each Sentence Verification trial presented a “True” and “False” response option in the top-left and top-right corners of the screen, respectively (Figure 5). Following countdown termination, a statement was displayed at the top-center of the screen, prompting participants to judge whether it was true or false by selecting the appropriate response option. Statement stimuli consisted of 21 simple declarative statements manipulated in truth value (true, false) and negation (non-negated, negated). Sentence negation was manipulated by adding “not” to statements (e.g., “giraffes are tall” is non-negated, while “giraffes are not tall” is negated). Truth value was manipulated by changing the adjective at the end of the sentence (e.g., “giraffes are not short” is true, while “giraffes are not tall” is false). Crossing these factors yielded four sentence conditions where each sentence could be a true or false statement in either negated or non-negated forms (see Figure 1 and Supplementary Table 4). Participants saw all four conditions of each statement, with the 84 resulting statements presented in a random order across trials.

#### 5.2.3 Photo Preference

Adapted from Koop and Johnson’s experiment [28] examining the mental dynamics of preferential choice, each Photo Preference trial presented the question “Which photo do you prefer?” centered at the top of the screen coincident with countdown initiation (Figure 5). Following countdown termination two images were then simultaneously displayed in the choice boxes to the upper left and upper right corners of the screen. As in Koop and Johnson [28], the International Affective Picture System (IAPS [29]) was used to develop a stimulus set of paired images using pleasantness ratings as an analog to photo preference, given equal levels of arousal [28]. We therefore selected 168 pictures from the IAPS, categorized as being high in pleasantness (pleasantness rating between 7 and 8), average in pleasantness (referred to as Med; pleasantness rating between 4.50 and 5.50) or low in pleasantness (pleasantness rating between 2 and 3). Images scoring greater than 6.15 in arousal were excluded. Selected pictures were then matched for arousal (difference *<* 0.30) and paired to create all pairwise combinations of High, Medium and Low. Pairs not matched in pleasantness (e.g., High*−*Med, High*−*Low, Med *−*Low) were counterbalanced for side of presentation, while pairs matched in pleasantness (e.g., High*−*High, Med *−*Med, Low *−*Low) appeared equally as often as the unmatched conditions when ignoring side of space (see Figure 1). This allowed for 14 presentations of each pleasantness pairing (7 of each unmatched pairing for each presentation side and 14 for matched pairings), for a total of 84 trials. Photo choice selections revealed a global preference for photos rated as more pleasant (*M* _*More PleasantSelected*_ = 78.3%), substantiating claims that preference is roughly analogous with pleasantness ratings [28]. As a result, the analysis included only trials containing a High pleasantness photo and in which the High photo was selected. Due to experimental error, half of participants completed a version of this task that did not counterbalance for side of presentation (i.e., High photos were always presented on the left). A separate ANOVA showed no significant difference between these groups for any measure, so both groups were included in the reported analysis where we collapsed across photo presentation side.

### 5.3 Data Treatment

#### 5.3.1 Operationalization of trajectory data

Raw movement data was resampled to 60 Hz, then filtered using a 10 Hz lowpass filter. Reach onset was defined as the first time the mouse cursor (Computer) or finger/thumb/stylus (Tablet and Smartphone) ascended to 5% of its peak velocity within the start button and after countdown had terminated. Should this velocity threshold not be achieved prior to leaving the start button, this threshold was iteratively reduced by 5% until a reach onset could be defined. Reach offset was similarly defined as the first time the mouse cursor (Computer) or finger/thumb/stylus (Tablet and Smartphone) velocity descended below a velocity threshold of 5% peak velocity while within one of the two choice option boxes, with this threshold iteratively increasing by 5% if necessary.

#### 5.3.2 Dependent Measures

For each trial, the following behavioural measures were obtained:

*Reaction time (seconds)*: time from countdown termination to reach onset.

*Movement time (seconds)*: time from reach onset to reach offset (choice selection).

*Trajectory curvature (MAD)*: Within each trial, the perpendicular distance of the observed trajectory relative to a straight line connecting the trajectory start and end positions was calculated for each data point. Maximum absolute deviation (MAD) reports the maximum of these perpendicular distances. Straight trajectories produce values approaching zero while those curving toward the center of the screen were assigned positive MAD values and those moving away from the center were assigned negative MAD values.

Within-participant and within-task z-scores were computed for each dependent measure (reaction time, movement time, trajectory curvature). This standardization of within-participant measures allows for between-task and between-participant comparisons while controlling for participant variability and individual reach patterns. All analyses were conducted on these standardized values. See Table 1 for reporting of raw and standardized measure values.

### 5.4 Data cleaning

Data cleaning processes were identical independent of device and were conducted using customized MATLAB scripts. Errors on each trial could be a combination of reaches with recording errors, reaches with insufficient data points (fewer than seven unique positions), reaches with reaction times greater than 0.1s, reaches with movement times *>* 3 SD above a participants mean movement time, and reaches with reaction times *>* 3 SD above a participants mean reaction time. For Numeric-Size Congruity and Sentence Verification tasks, incorrect trials were also removed from analysis. As these tasks previously demonstrated very high levels of accuracy [13, 17], incorrect responses were considered to arise from participant error, with sustained performance errors indicating participant unreliability. The average percentage of total participant trials falling within each of these error categories are reported in Supplementary Table 5. A participant was excluded from analysis if, after data cleaning, they failed to have at least four trials in each condition of analysis as reported per task. In total, participants whose data was included for analysis had a mean of 95.6% usable trials for analysis (Range: 83.7%*−*98.4%).

### 5.4 Analysis

The main objective of this analysis was to determine whether task-specific decision-difficulty effects (as expected by previous studies, e.g., [17, 28, 13, 31]) were replicated and whether these effects were consistent despite differences in testing device. To that end, analysis proceeded in three primary stages: 1) a-priori comparisons to determine replication of antecedent results, 2) within-task, between-device omnibus analysis of variances (ANOVAs) to determine any effects or interactions arising due to device differences, and 3) between-device ANOVA to determine whether there are correlational relationships between measures of decision-difficulty and if these remain consistent across device.

#### 5.4.1 A-Priori Comparison Procedure

To determine replication of the previous task-specific difficulty effects, a subset of trial conditions were selected to represent low and high difficulty decisions within each task (see Figure 1). For the Numeric-Size Congruity task, decision-difficulty followed size-congruity, with trials incongruent in numerical and physical size categorized as high in decision-difficulty, while congruent trials were categorized as low in decision-difficulty [17]. For the Sentence Verification task, decision-difficulty varied according to negation, with true statements the greatest negation-driven effects [13]. The current study therefore categorized true negated trials as representative of a high difficulty decision, and true non-negated trials as having low decision-difficulty. Finally, decision-difficulty in the Photo Preference task was driven by the similarity in pleasantness between photos [28]. The current study places trials comparing two photos high in pleasantness in the high decision-difficulty category, and trials comparing a photo high in pleasantness and one low in pleasantness in the low decision-difficulty category. Within each task, mean standardized reaction time, movement time and trajectory curvature scores for low and high decision-difficulty trials were compared using a paired t-test. As these were a-priori tests based on replicating known effects, significance was set to *p≤*.05 with no correction for multiple comparisons.

#### 5.4.2 Within-task ANOVA Procedure

Mean standardized reaction time, movement time and maximum absolute deviation measures were separately submitted to mixed-model ANOVAs, with within-subject factors determined by individual tasks design and between-subject factors of device (computer, tablet, smartphone, see section 2 - Results). All multi-way mixed- and RM-ANOVAs were family-wise error corrected using a sequential Bonferroni procedure [11], and all repeated-measures main effects and interactions were Greenhouse-Geiser corrected to protect against violations of sphericity. The primary objective of this series of tests was to look for device differences. As a result, here we focus only on main effects or interactions involving Device. Full results outside this explicit objective can be found in Supplementary Materials 1, including results that support the a-priori tests of decision-difficulty. Interactions involving Device first collapsed over factors that did not interact, then were followed up by separating by the factor(s) other than Device. Significant (simple) main effects of Device were explored with all possible pairwise comparisons which were Bonferroni corrected with significance set at a corrected *p≤*.01.

#### 5.4.3 Between-task ANOVA Procedure

To explore the relationship between measures of decision-difficulty, a Pearson’s correlation coefficient (*r*) was calculated between each pair of measures (*r* _*MAD,MT*_, *r* _*MAD,RT*_ and *r* _*MT,RT*_) indicating the direction and strength of the relationship across trials for each participant within each condition, task, and device. Mean correlation coefficients were then submitted to a mixed-model ANOVA with Correlation-type and Task as within-subjects factors and Device as a between-subjects factor. Corrections and follow-up procedures were then conducted as described above, except here we were most interested in the pairwise comparisons between Task.

## Supporting information

Complete Supplementary Materials

## Acknowledgments

This study was funded by an NSERC Discovery Grant to CSC, and by an NSERC CGS-M and MITACS Accelerate International award to AOZ.

## References

[1] Herman Aguinis, Isabel Villamor, and Ravi S Ramani. “MTurk research: Review and recommendations”. In: Journal of Management 47.4 (2021), pp. 823–837.

[2] Alexander Anwyl-Irvine et al. “Realistic precision and accuracy of online experiment platforms, web browsers, and devices”. In: Behavior research methods 53.4 (2021), pp. 1407–1425.

[3] Alexander L Anwyl-Irvine et al. “Gorilla in our midst: An online behavioral experiment builder”. In: Behavior research methods 52.1 (2020), pp. 388–407.

[4] Pavlo Bazilinskyy and Joost C. F. de Winter. “Crowdsourced Measurement of Reaction Times to Audiovisual Stimuli With Various Degrees of Asynchrony”. In: Human Factors (2018). doi: 10.1177/0018720818787126.

[5] Daniel Bernoulli. “Exposition of a New Theory on the Measurement of Risk”. In: Econometrica (1954). doi: 10.2307/1909829.

[6] Diana Burk et al. “Motor effort alters changes of mind in sensorimotor decision making.” In: PLOS ONE (2014). doi: 10.1371/journal.pone.0092681.

[7] Craig S. Chapman et al. “Reaching for the unknown: multiple target encoding and real-time decisionmaking in a rapid reach task”. In: Cognition (2010). doi: 10.1016/j.cognition.2010.04.008.

[8] Craig S. Chapman et al. “Short-term motor plasticity revealed in a visuomotor decision-making task.” In: Behavioural Brain Research (2010). doi: 10.1016/j.bbr.2010.05.012.

[9] Paul Cisek and John F. Kalaska. “Neural Mechanisms for Interacting with a World Full of Action Choices”. In: Annual Review of Neuroscience (2010). doi: 10.1146/annurev.neuro.051508.135409.

[10] Ignasi Cos, Farid Medleg, and Paul Cisek. “The modulatory influence of end-point controllability on decisions between actions.” In: Journal of Neurophysiology (2012). doi: 10.1152/jn.00081.2012.

[11] Angélique O. J. Cramer et al. “Hidden multiplicity in exploratory multiway ANOVA: Prevalence and remedies.” In: Psychonomic Bulletin & Review (2016). doi: 10.3758/s13423-015-0913-5.

[12] Helen A. Cunningham and Robert B. Welch. “Multiple concurrent visual-motor mappings: implications for models of adaptation.” In: Journal of Experimental Psychology: Human Perception and Performance (1994). doi: 10.1037/0096-1523.20.5.987.

[13] Rick Dale and Nicholas D. Duran. “The Cognitive Dynamics of Negated Sentence Verification”. In: Cognitive Science (2011). doi: 10.1111/j.1551-6709.2010.01164.x.

[14] Dror Dotan et al. “On-line confidence monitoring during decision making”. In: Cognition (2018). doi: 10.1016/j.cognition.2017.11.001.

[15] Dror Dotan et al. “Track It to Crack It: Dissecting Processing Stages with Finger Tracking.” In: Trends in Cognitive Sciences (2019). doi: 10.1016/j.tics.2019.10.002.

[16] Christopher D. Erb, Jeff Moher, and Stuart Marcovitch. “Attentional capture in goal-directed action during childhood, adolescence, and early adulthood.” In: Journal of Experimental Child Psychology (2022). doi: 10.1016/j.jecp.2021.105273.

[17] Thomas J. Faulkenberry et al. “Response trajectories capture the continuous dynamics of the size congruity effect”. In: Acta Psychologica (2016). doi: 10.1016/j.actpsy.2015.11.010.

[18] Holger Finger et al. “LabVanced: a unified JavaScript framework for online studies”. In: International conference on computational social science (cologne). 2017.

[19] Michael C. Frank et al. “Using Tablets to Collect Data From Young Children”. In: Journal of Cognition and Development (2016). doi: 10.1080/15248372.2015.1061528.

[20] Jonathan B. Freeman. “Doing Psychological Science by Hand”. In: Current Directions in Psychological Science (2018). doi: 10.1177/0963721417746793.

[21] Jason P. Gallivan and Craig S. Chapman. “Three-dimensional reach trajectories as a probe of realtime decision-making between multiple competing targets.” In: Frontiers in Neuroscience (2014). doi: 10.3389/fnins.2014.00215.

[22] Jason P. Gallivan et al. “Decision-making in sensorimotor control.” In: Nature Reviews Neuroscience (2018). doi: 10.1038/s41583-018-0045-9.

[23] Claude Ghez et al. “Discrete and continuous planning of hand movements and isometric force trajectories”. In: Experimental Brain Research (1997). doi: 10.1007/pl00005692.

[24] Claude Ghez et al. “Roles of proprioceptive input in the programming of arm trajectories.” In: Cold Spring Harbor Symposia on Quantitative Biology (1990). doi: 10.1101/sqb.1990.055.01.079.

[25] Adrian M. Haith, David M. Huberdeau, and John W. Krakauer. “Hedging your bets: intermediate movements as optimal behavior in the context of an incomplete decision.” In: PLOS Computational Biology (2015). doi: 10.1371/journal.pcbi.1004171.

[26] Eric Hehman, Ryan M. Stolier, and Jonathan B. Freeman. “Advanced mouse-tracking analytic techniques for enhancing psychological science”. In: Group Processes & Intergroup Relations (2015). doi: 10.1177/1368430214538325.

[27] Mel W. Khaw, Paul W. Glimcher, and Kenway Louie. “Normalized value coding explains dynamic adaptation in the human valuation process.” In: Proceedings of the National Academy of Sciences of the United States of America (2017). doi: 10.1073/pnas.1715293114.

[28] Gregory J. Koop and Joseph G. Johnson. “The response dynamics of preferential choice.” In: Cognitive Psychology (2013). doi: 10.1016/j.cogpsych.2013.09.001.

[29] Pj Lang. “International affective picture system (IAPS) : affective ratings of pictures and instruction manual”. In: CTIT technical reports series (2005). doi: null.

[30] Joshua de Leeuw. “JsPsych: a JavaScript library for creating behavioral experiments in a Web browser.” In: Behavior Research Methods (2015). doi: 10.3758/s13428-014-0458-y.

[31] Mora Maldonado, Ewan Dunbar, and Emmanuel Chemla. “Mouse tracking as a window into decision making.” In: Behavior Research Methods (2019). doi: 10.3758/s13428-018-01194-x.

[32] Gregory McCarthy and Emanuel Donchin. “A metric for thought: a comparison of P300 latency and reaction time”. In: Science (1981). doi: 10.1126/science.7444452.

[33] Jeff Moher and Joo-Hyun Song. “Perceptual decision processes flexibly adapt to avoid change-of-mind motor costs.” In: Journal of Vision (2014). doi: 10.1167/14.8.1.

[34] Camillo Padoa-Schioppa and John A. Assad. “Neurons in the orbitofrontal cortex encode economic value”. In: Nature (2006). doi: 10.1038/nature04676.

[35] Stefan Palan and Christian Schitter. “Prolific.ac—A subject pool for online experiments”. In: Journal of Behavioral and Experimental Finance (2017). doi: 10.1016/j.jbef.2017.12.004.

[36] John Palmer, Alexander C. Huk, and Michael N. Shadlen. “The effect of stimulus strength on the speed and accuracy of a perceptual decision”. In: Journal of Vision (2005). doi: 10.1167/5.5.1.

[37] Eliza Passell et al. “Cognitive test scores vary with choice of personal digital device”. In: Behavior Research Methods (2021). doi: 10.3758/s13428-021-01597-3.

[38] John W. Payne. “Task complexity and contingent processing in decision making: An information search and protocol analysis”. In: Organizational Behavior and Human Performance (1976). doi: 10.1016/0030-5073(76)90022-2.

[39] Thomas Pronk et al. “Can we measure individual differences in cognitive measures reliably via smartphones? A comparison of the flanker effect across device types and samples”. In: Behavior Research Methods (2022), pp. 1–12.

[40] Antonio Rangel and Todd A. Hare. “Neural computations associated with goal-directed choice”. In: Current Opinion in Neurobiology (2010). doi: 10.1016/j.conb.2010.03.001.

[41] Michael Schulte-Mecklenbeck et al. “Process-tracing methods in decision making: on growing up in the 70s”. In: Current Directions in Psychological Science (2017). doi: 10.1177/0963721417708229.

[42] Kilian Semmelmann et al. “U Can Touch This: How Tablets Can Be Used to Study Cognitive Development”. In: Frontiers in Psychology (2016). doi: 10.3389/fpsyg.2016.01021.

[43] Britne A. Shabbott and Robert L. Sainburg. “Learning a visuomotor rotation: simultaneous visual and proprioceptive information is crucial for visuomotor remapping”. In: Experimental Brain Research (2010). doi: 10.1007/s00221-010-2209-3.

[44] Paul E. Stillman, Xi Shen, and Melissa J. Ferguson. “How Mouse-tracking Can Advance Social Cognitive Theory”. In: Trends in Cognitive Sciences (2018). doi: 10.1016/j.tics.2018.03.012.

[45] Paul E. Stillman et al. “Using dynamic monitoring of choices to predict and understand risk preferences.” In: Proceedings of the National Academy of Sciences of the United States of America (2020). doi: 10.1073/pnas.2010056117.

[46] Fritz Strack, Leonard L. Martin, and Sabine Stepper. “Inhibiting and facilitating conditions of the human smile: A nonobtrusive test of the facial feedback hypothesis.” In: Journal of Personality and Social Psychology (1988). doi: 10.1037/0022-3514.54.5.768.

[47] Nicolette Sullivan et al. “Dietary self-control is related to the speed with which attributes of healthfulness and tastiness are processed”. In: Psychological science 26.2 (2015), pp. 122–134.

[48] Grace Truong et al. “Mine in motion: how physical actions impact the psychological sense of object ownership”. In: Journal of Experimental Psychology: Human Perception and Performance (2016). doi: 10.1037/xhp0000142.

[49] Kunlin Wei et al. “Computer Use Changes Generalization of Movement Learning”. In: Current Biology (2014). doi: 10.1016/j.cub.2013.11.012.

[50] Nathan J. Wispinski, Jason P. Gallivan, and Craig S. Chapman. “Models, movements, and minds: bridging the gap between decision making and action”. In: Annals of the New York Academy of Sciences (2020). doi: 10.1111/nyas.13973.

[51] Nathan J. Wispinski et al. “Reaching reveals that best-versus-rest processing contributes to biased decision making.” In: Acta Psychologica (2017). doi: 10.1016/j.actpsy.2017.03.006.

[52] Aaron L. Wong and Adrian M. Haith. “Motor planning flexibly optimizes performance under uncertainty about task goals.” In: Nature Communications (2017). doi: 10.1038/ncomms14624.

[53] Kenji Yamamoto et al. “Rapid and long-lasting plasticity of input-output mapping.” In: Journal of Neurophysiology (2006). doi: 10.1152/jn.00209.2006.

