## Supplementary material for "Continuous Measures of Decision-Difficulty Captured Remotely: I. Mouse-tracking sensitivity extends to tablets and smartphones": Complete Supplementary Materials

The complete Supplementary Materials includes the following items:

**Supplementary Note:**

1) Trajectory visualization details

**Supplementary Discussions:**

1) Confirmation of a-priori analysis results from omnibus ANOVA analyses
2) Difficulty effects driven by numerical pairing in Numeric-Size Congruity task
3) Further support of right-hand bias during touch-device use

**Supplementary Tables:**

1) Table of demographic survey responses
2) Table of device specifications reported by computer users
3) Table of device specifications reported by touch-device users
4) Table of Sentence Verification statement stimuli
5) Table of non-mutually exclusive errors constituting trial removal by device

**Supplementary Materials References**

---

**Supplementary Note 1: Trajectory visualization details**

Trajectories were extracted as described in the main manuscript. For visualizations, the following steps were taken. First, all trajectories were space normalized to the y-dimension (up/down on screen) using functional data analysis (see Gallivan & Chapman, 2014). Then, all trajectories ending on the right side were horizontally flipped so that they ended on the left (thus all trajectories ended left). Then, we plot each participant's space-normalized mean trajectory across all trials for the given high/low decision difficulty comparison. These individual means are shown as thin gray lines. To show the high/low decision-difficult grand average trajectories in a way that more authentically maps to our measure of standardized MAD we represented each space-normalized trajectory from every individual by its perpendicular distances relative to its own straight line (connecting its start to its end). We recorded the parameters of each straight line and then we z-scored *all* of these perpendicular distances (e.g., each point on each trajectory) by the same distribution (e.g., divide by the standard deviation) that defined the z-score of the MAD values for each participant across all of their trials. This allowed us, for each participant, to get a condition mean (e.g., all the low-difficulty trials) represented as the average space-normalized straight line and average set of z-scored perpendicular distances along that line. To convert these mean z-score distances back into a space that maps to the pixel coordinates we multiplied them by the same standard deviation that had been used to z-score each raw data point (e.g., the one derived from the MAD distribution for each participant). These standardized pixel distances were then added to the mean straight line for each individual and each condition to give a mean standardized trajectory for that condition for that individual. The thick condition colored lines in the plot and inset are the average of these standardized trajectories across all participants. To calculate the error bars shown in the inset plots, we used a similar procedure. At the step where we calculated the average set of z-scored perpendicular distances along the straight line for each individual for a particular condition, we also calculated the standard error of these perpendicular distances. These were converted from z-scores into pixel space using the same standard deviation multiplication as was used for the mean z-score distances. That meant, in addition to a mean standardized set of pixel distances per individual and condition, we also had an estimate of the standard error, in pixels, over that mean set of distances (e.g., the trajectory). What we visualize as error bars on the inset plots is the mean trajectory plus and minus the mean of this standard error averaged across all individuals. It is best thought of as an average of within-subjects standard error, but here, it has gone through a conversion to more accurately map to the z-score MAD values.

**Supplementary Discussion 1: Confirmation of a-priori analysis results from omnibus ANOVA analyses**

To assess any effects due to the device used, in the main text we submitted mean standardized reaction times, movement times and trajectory curvature scores for each task to a mixed-model ANOVA (see "*Within-task ANOVA Procedure*" for further details), but only looked for main effects or interactions involving the Device factor and explored any simple main effects with pairwise comparisons between levels of Device. Here, we report all task-specific results falling outside that narrow scope, and discuss their confirmation of our a-priori t-test analyses. Together, these results support our replication of previous task-specific decision-difficulty effects across all three devices assessed as presented in "*Results 1: Tablets and smartphones measure decision difficulty as well as computer mouse-tracking during reach-decision tasks*".

**1.1. Numeric-Size Congruity**

***Reaction Time Results.*** Beyond those effects reported in the main text, the 4-factor Number Pairs x Congruity x Presentation Side x Device mixed-model ANOVA for Numeric-Size Congruity reaction time data revealed main effects of Number Pairs ($F(2,237) = 77.15$, $p = 2.46e-27$, $\eta 2 = .061$), Congruity ($F(1,237) = 137.57$, $p = 2.36e-25$, $\eta 2 = .050$), as well as a two-way Number Pairs x Congruity interaction ($F(2,237) = 9.09$, $p = 1.34e-4$, $\eta 2 = .005$). Simple main effect 1-factor RM-ANOVAs assessing Congruity at each level of Number Pairs revealed significant main effects of Congruity at all three levels (1 – 2: $F(1,238) = 79.39$, $p = 1.38e-16$; 2 – 8: $F(1,238) = 17.39$, $p = 4.28e-5$; 8 – 9: $F(1,238) = 75.55$, $p = 5.97e-16$). In line with results observed in our a-priori analysis, Incongruent pairings showed greater standardized reaction time scores compared to Congruent pairings at all three levels (1v2: $M_{Incongruent-Congruent} = 0.23$, $t = 8.64$, $p = 5.79e-16$, $d = 0.57$; 2v8: $M_{Incongruent-Congruent} = 0.11$, $t = 3.99$, $p = .001$, $d = 0.26$; 8v9: $M_{Incongruent-Congruent} = 0.25$, $t = 9.29$, $p = 2.82e-18$, $d = 0.61$) indicating significant difference in decision difficulty as a function of Congruity wherein Incongruent trials are more difficult than Congruent trials. These effects were also reliably modulated by the numeric values of the digits being compared, 8v9 digit pairings displayed the greatest overall decision difficulty and greatest congruity effects (see "*Supplemental Discussion 2*" for discussion of the basis of this modulation). Notably, the absence of a Device effect or Interaction is evidence for the consistency for decision-difficulty to shape reaction times for this task, across devices.

***Movement Time Results.*** The 4-factor Number Pairs x Congruity x Presentation Side x Device mixed-model ANOVA for Numeric-Size Congruity movement time data revealed main effects of Number Pairs ($F(2,237) = 236.54$, $p = 2.36e-69$, $\eta2 = .129$), Congruity ($F(1,237) = 186.48$, $p = 1.04e-31$, $\eta2 = .052$), as well as a two-way Number Pairs x Congruity interaction ($F(2,237) = 22.12$, $p =9.81e-10$, $\eta2 = .012$). Subsequent simple main effect 1-factor RM-ANOVAs assessing Congruity at each level of Number Pairs revealed significant main effects of Congruity at all three levels (1v2: $F(1,238) = 47.70$, $p = 4.54e-11$; 2v8: $F(1,238) = 18.08$, $p = 3.05e-5$; 8v9: $F(1,238) = 136.39$, $p = 3.44e-25$). As with the reaction time scores, Incongruent pairs showed greater standardized movement time scores compared to Congruent pairs in all cases (1v2: $M_{Incongruent-Congruent} = 0.20$, $t = 6.95$, $p = 1.22e-10$, $d = 0.47$; 2v8: $M_{Incongruent-Congruent} = 0.11$, $t = 3.69$, $p = .004$, $d = 0.25$; 8v9: $M_{Incongruent-Congruent} = 0.37$, $t = 12.97$, $p = 4.34e-33$, $d = 1.10$). In line with our a-priori analysis results and expected number-pair driven results (see "*Supplemental Discussion 2*"), this interaction again suggests greater relative decision difficulty for Incongruent trials, with 8v9 pairs showing greater overall difficulty (i.e., longest movement times) and congruity effects. Of note, the absence of an interaction between these two interacting factors and Device suggest instead that this effect does not differ across the computer-, tablet-, and smartphone-based testing.

***Trajectory Results.*** In addition to the effects reported in the main text, the 4-factor Number Pairs x Congruity x Presentation Side x Device mixed-model ANOVA assessing Numeric-Size Congruity MAD trajectory data revealed main effects of Number Pairs ($F(2,237) = 235.25$, $p = 9.46e-67$, $\eta 2 = .077$), Congruity ($F(1,237) = 395.13$, $p = 2.12e-52$, $\eta 2 = .074$), and a two-way Number Pairs x Congruity interaction ($F(2,237) =99.46$, $p = 9.17e-34$, $\eta 2 = .030$).

As with reaction time and movement time analyses, three simple main effect 1-factor RM-ANOVAs were used to compare Congruity at each level of Number Pairs, and revealed significant main effects of Congruity at all three levels (1v2: $F(1,238) = 54.96$, $p = 2.17e-12$; 2v8: $F(1,238) = 81.52$, $p = 6.18e-17$; 8v9: $F(1,238) = 335.15$, $p = 2.96e-47$). In all cases, Incongruent pairs again showed greater standardized MAD scores compared to Congruent pairs, with the greatest Congruity effect revealed for 8v9 pairs compared to 1v2 and 2v8 pairs (1v2: $M_{Incongruent-Congruent} = 0.17$, $t = 6.17$, $p = 1.69e-8$, $d = 0.32$; 2v8: $M_{Incongruent-Congruent} = 0.20$, $t = 7.43$, $p = 4.71e-12$, $d = 0.39$; 8v9: $M_{Incongruent-Congruent} = 0.63$, $t = 23.37$, $p = 1.27e-88$, $d = 1.23$). These results align with the decision difficulty effects revealed within reaction time and movement time, ultimately replicating predicted results. No interaction between our effect of interest (numeric distanced-modulated Congruity and Device) was revealed. As with our movement time analysis, this again suggests that these effects do not differ across the computer-, tablet-, and smartphone-based testing.

### 1.2. Sentence Verification

***Movement Time Results.*** Not presented in the main text, the 3-factor Truth Value x Negation x Device mixed-model ANOVA for Sentence Verification standardized movement time scores revealed main effects for Truth Value ($F(1,237) = 16.97$, $p = 5.25e-5$, $\eta 2 = .012$), Negation ($F(1,237) = 615.84$, $p = 7.66e-68$, $\eta 2 = .43$), and a two-way interaction between these factors ($F(1,237) = 192.00$, $p = 2.23e-32$, $\eta 2 = .086$. Tests conducted to follow up the Truth Value x Negation interaction revealed a main effect of Negation when statements were True ($F(1,238) = 618.93$, $p = 4.99e-68$) as well as when they were False ($F(1,238) = 153.51$, $p = 1.63e-27$). In both cases, Negated statements showed greater standardized movement time scores compared to Non-negated statements (True: $M_{Negated-Non-negated} = 0.74$, $t = 28.01$, $p = 3.22e-100$, $d = 2.80$; False: $M_{Negated-Non-negated} = 0.28$, $t = 10.60$, $p = 7.82e-23$, $d = 1.07$) In line with our a-prior analysis results, here we see an increase in decision difficulty expressed by movement time during the verification of negated statements compared to the verification of non-negated statements (with greater differences between negation conditions for True statements than False statements). These interconnected truth value and negation effects do not interact with Device, suggesting consistent replication of these effects across computer-, tablet- and smartphone-based testing.

***Trajectory Curvature Results.*** As with movement time, the 3-factor Truth Value x Negation x Device mixed-model ANOVA for Sentence Verification standardized MAD scores revealed main effects for Truth Value ($F(1,237) = 30.83$, $p = 7.52e-8$, $\eta 2 = .075$), Negation ($F(1,237) = 125.04$, $p = 1.37e-23$, $\eta 2 = .050$), and a two-way interaction between these factors ($F(1,237) = 96.53$, $p = 2.48e-19$, $\eta 2 = .034$). A simple main effect 1-factor RM-ANOVA revealed a main effect of

Negation when statements were True ($F(1,238) = 156.14$, $p = 7.35e-28$), but not when they were False. As in our a-priori analysis, those True statements that were Negated showed greater standardized MAD values compared to those that were Non-negated ($M_{Negated-Non-negated} = 0.39$, $t = 14.88$, $p = 1.49e-40$, $d = 0.93$). The results again suggest negation-driven decision difficulty effects which are diminished when statements are False compared to when they are True. Device was not shown to modulate this interaction as it is reflected in trajectory curvature.

**1.3. Photo Preference**

***Movement Time Results.*** A 2-factor Valence Pairing x Device mixed-model ANOVA applied to Photo Preference standardized movement time scores revealed only a main effect of Valence Pairing ($F(2,237) = 24.27$, $p =1.52e-10$, $\eta 2 = .060$), with pairwise comparisons showing High – High to have significantly greater movement times compared to High – Med pairings ($M_{HH-HM} = 0.093$, $t = 3.63$, $p = 9.51e-4$, $d = 0.33$) and High – Low pairings ($M_{HH-HL} = 0.18$, $t = 6.97$, $p = 3.35e-11$, $d = 0.63$). Paired with the absence of any interaction with Device, these results suggest that decision difficulty does indeed increase as similarity in photo pleasantness increases and that this effect does not differ as a result of testing device, supporting a-priori analysis results.

***Trajectory Curvature Results.*** As with the trajectory curvature analysis, the 3-factor Valence Pairing x Device mixed-model ANOVA applied to Photo Preference standardized MAD scores revealed only a main effect of Valence Pairing ($F(2,237) = 35.08$, $p = 5.36e-14$, $\eta 2 = .086$). As with movement time, High – High were shown to have significantly greater movement times compared to High – Med pairings ($M_{HH-HM} = 0.19$, $t = 6.88$, $p = 5.81e-11$, $d = 0.64$) and High – Low pairings ($M_{HH-HL} = 0.21$, $t = 7.58$, $p = 5.71e-13$, $d = 0.70$). These results again support a-priori results indicating an increase in decision difficulty as photo pleasantness increased in similarity. This analysis also did not reveal any interaction or main effect of Device, serving as an additional indicator that trajectory-curvature expressed decision difficulty is consistent independent of testing device used.

**Supplementary Discussion 2: Difficulty effects driven by numerical pairing in Numeric-Size Congruity task**

For the purpose of our replication of difficulty-driven effects across devices, we focussed on size-congruity as the main driver of decision difficulty in the Numeric-Size Congruity task. However, this task presented a second dimension of difficulty in the number pairs being tested. Briefly, the numerical pair with the largest numbers (8v9) leads to a harder decision than numerical pairs with one or more of the smaller numbers (1v2 and 2v8). This is due to two properties that impact numerical judgements: 1) the distance between two numbers where numbers spaced further apart on the number line are easier to distinguish (Schwarz & Ischebeck, 2003; Santens, Goossens & Verguts, 2011) and 2) the relative size of the numerical difference where differences of similar size are harder to differentiate  (Moyer & Landauer, 1967) particularly as the constituent numbers get larger ( as we are exposed to larger numbers less often, Dehaene and Mehler, 1992). While our a-priori analysis allowed us to focus our assessment on task decision difficulty effects driven by congruity, our mixed-model omnibus ANOVA allowed the assessment of decision difficulty effects using at the level of number pair. The interaction between this level of decision difficulty and device are presented in "*Results 2.1: Measure sensitivity pre-movement"*. Non-device related decision difficulty results related to this effect are discussed further in "*Supplementary Discussion 1"*.

**Supplementary Discussion 3: Further support of right-hand bias during touch-device use**

A right-hand bias was also present in Numeric-Size Congruity movement time results. As with the trajectory curvature results for this task reported in the main text ("*Results 2.2: Measure sensitivity during-movement"*), our movement time analysis demonstrated a number pair Presentation Side x Device interaction ($F(1,237) = 18.07$, $p = 4.96e-81$, $\eta 2= .019$), and main effects of Device for both Left ($F2,237) = 18.656$, $p = 2.960e-8$) and Right reaches ($F(2,237) = 17.872$, $p = 5.842e-8$) driven by significant differences between Computers and Tablets (Left: $M_{Computer-Tablet} = -0.15$, $t = 5.50$, $p = 1.42e-6$, $d = 0.36$; Right: $M_{Computer-Tablet} = 0.15$, $t = 5.53$, $p = 1.21e-6$, $d = 0.36$) and Computers and Smartphones (Left: $M_{Computer-Smartphone} = -0.13$, $t = 4.59$, $p = 1.05e-4$, $d = 0.30$, Right:$M_{Computer-Smartphone} = 0.13$, $t = 4.72$, $p = 6.01e-5$, $d = 0.31$) as a result of the touch-device movement times demonstrating faster movement times during rightward reaches (Tablet: $M_{Right-Left} = 0.30$, $t = 7.48$, $p = 2.04e-11$, $d = 0.70$; Smartphone: $M_{Right-Left} = 0.25$, $t = 6.26$, $p = 2.74e-8$, $d = 0.59$) while computers did not. While the preferential processing of stimuli presented on the right can manifest as right-hand biases in trajectory curvature (as seen in the results reported in the main text), this can also be paired with faster movement times during rightward reaches (Chapman et al., 2010b; Gallivan & Chapman, 2014). The movement time directional effects presented here therefore provided further support for our observations of right-hand bais during touch-device use compared to computer-use.

**Supplementary Table 1.** Table of demographic survey responses

| Survey Questions | Response Options | Disclosed Responses (count) | | |
|---|---|---|---|---|
| | | Computer | Tablet | Smartphone |
| | | *N = 83* | *N = 79* | *N = 80* |
| What gender do you identify as? | Male | 56 | 51 | 52 |
| | Female | 25 | 27 | 25 |
| | Prefer not to say | 2 | | 1 |
| | Other *(custom response)* | | 1 | |
| What is your age? | *(custom response)* | range [21-65] | range [23-58] | range [24-57] |
| What hand do you typically write with (i.e. your dominant hand)? | Right | 76 | 67 | 71 |
| | Left | 6 | 12 | 7 |
| Do you have normal or corrected to normal vision? (vision correction may be via surgery, glasses, contacts, etc.) | Yes | 79 | 77 | 73 |
| | No | 4 | 2 | 5 |
| Is English your first language? | Yes | 80 | 77 | 76 |
| | No | 2 | 2 | 2 |
| If English is not your first language, how old were you when you learned English? | *(custom response)* | 4,23 | 3,4 | 3.5 |
| How would you rate your English reading comprehension skills? (1-5) | 1 | | | 1 |
| | 2 | | | |
| | 3 | | | |
| | 4 | 2 | 2 | 3 |
| | 5 | 81 | 77 | 74 |
| How many hours a week do you play video games? | 0 | 2 | 2 | 2 |
| | Less than 1 | 6 | 6 | 3 |
| | 1 - 3 | 16 | 13 | 15 |
| | 4 - 6 | 16 | 14 | 14 |
| | 7 - 10 | 12 | 17 | 14 |
| | 11 - 15 | 7 | 6 | 10 |
| | 16 - 20 | 11 | 5 | 8 |
| | 20+ | 13 | 14 | 12 |
| How many hours a week do you participate in activities requiring coordinated hand - eye movement? (e.g. instrument playing, catching and throwing, swinging, dribbling, etc.) | 0 | 8 | 1 | 4 |
| | Less than 1 | 21 | 14 | 10 |
| | 1 - 3 | 18 | 26 | 24 |
| | 4 - 6 | 12 | 14 | 11 |
| | 7 - 10 | 4 | 7 | 11 |
| | 11 - 15 | 6 | 3 | 4 |
| | 16 - 20 | 2 | 5 | 3 |
| | 20+ | 12 | 7 | 11 |

**Supplementary Table 2.** Table of device specifications reported by computer users

| Survey Questions | Selected Response Options (count) | |
|---|---|---|
| **Desktop Computer** (n = 69) | | |
| What brand of desktop computer are you using? | Apple (1) | Lenovo (1) |
| | Acer (1) | Microsoft (5) |
| | Asus (9) | I don't know (3) |
| | HP (13) | Other *(custom response)* (25*) |
| | Dell (9) | |
| What brand is your monitor? | Apple (3) | HP (7) |
| | Acer (11) | Lenovo (2) |
| | Asus (10) | I don't know (2) |
| | Dell (13) | Other *(custom response)* (21**) |
| What size is your monitor? | 22 inches (10) | 34 inches (4) |
| | 24 inches (28) | I don't know (2) |
| | 27 inches (19) | Other *(custom response)* (6***) |
| What operating system does your computer use? | Windows (64) | Linux (1) |
| | Mac OS (2) | I don't know (1) |
| | Chrome OS (1) | |
| How are you choosing to interact with your computer? | Wired mouse (52) | Wireless mouse (17) |
| **Laptop Computer** (n = 14) | | |
| What brand of laptop computer are you using? | Acer (3) | HP (6) |
| | Dell (4) | Other *(custom response)* (1 †) |
| What model/series of laptop from the brand are you using? | *(custom response)* †† | |
| What is your laptop's screen size? (round to nearest option if necessary) | 11 inches (1) | 16 inches (1) |
| | 14 inches (2) | 17 inches (5) |
| | 15 inches (5) | |
| What operating system does your laptop use? | Windows (13) | Chrome OS (1) |
| How are you choosing to interact with your laptop? | Wired mouse (4) | Wireless mouse (10) |

Disclosed custom responses (note, only unique responses are reported here):

* Custom build, CyberPowerPC, MAINGEAR, iBUYPOWER, CORSAIR

** AOC, BenQ, LG, Panasonic (television), Samsung, ViewSonic, Planar

*** 16 inches, 20 inches, 30 inches, 32 inches, 42 inches

† RCA

†† Acer Chromebook 15, Acer Nitro 5, Acer Aspire 3, Dell Inpsiron 7300, HP ZBook, Dell Inpsiron 15 5570, Dell Inspiron 15 5000, Dell XPS 15, Dell Vostro 3550, HP EliteBook, HP Pavilon, HP Spectre x360, RCA Cambio Windows 2-in-1 Tablet/Laptop

**Supplementary Table 3.** Table of device specifications reported by touch-device users

| Survey Questions | Selected Response Options (count) | |
|---|---|---|
| **Tablet** (n = 79) | | |
| What brand of tablet are you using? | Samsung (39)<br>Google (6)<br>Amazon (25) | I don't know (2)<br>Other *(custom response)* (7*) |
| What is the make and model/series of the tablet you are using? | *(custom response)*** |  |
| What is your tablet's screen size? (round to the nearest option if necessary) | 7 inches (15)<br>8 inches (23)<br>9 inches (4)<br>10 inches (28) | 11 inches (4)<br>I don't know (4)<br>Other *(custom response) (1***)* |
| How are you choosing to interact with your tablet? | Stylus on touchscreen (5) | Finger/thumb on touchscreen (74) |
| **Smartphone** (n = 78) | | |
| What brand of smartphone are you using? | Samsung (45)<br>Google (4)<br>LG (10)<br>Nokia (2) | Motorola (6)<br>Huawei (2)<br>Other *(custom response)* (9 †) |
| What is the make and model/series of the smartphone you are using? | *(custom response)* †† | |
| How are you choosing to interact with your smartphone? | Finger on touchscreen (43)<br>Thumb on touchscreen (34) | Stylus on touchscreen (1) |

Disclosed custom responses (note, only unique responses are reported here):

* Acer, Lenovo, RCA, TECNO

** Acer Iconia One 7, Amazon Fire, Amazon Fire HD, Amazon Fire 7, Amazon Fire 8, Amazon Fire 8 HD, Amazon Fire 10, Amazon Fire 10 HD, Amazon Kindle, Amazon Kindle Fire, Amazon Kindle Fire 5, Amazon Kindle Fire HD, Fusion, Google Nexus 7, Google Pixel Slate, Lenovo M10 Plus, Lenovo Tab 4, ONN Tablet, RCA Galileo, Samsung Galaxy Tab 2, Samsung Galaxy Tab 4, Samsung Galaxy Tab A, Samsung Galaxy Tab A 10.1, Samsung Galaxy Tab A6, Samsung Galaxy Tab A7, Samsung Galaxy Tab A8, Samsung Galaxy Tab E, Samsung Galaxy Tab S10, Samsung Galaxy Tab S2, Samsung Galaxy Tab S3, Samsung Galaxy Tab S6, Samsung Galaxy Tab S6 Lite, Samsung Galaxy Tab S7, TECNO DroiPad, I don't know

*** 12.4

† Asus, BLU, Realme, TECNO, OnePlus, Xiaomi

†† Asus ROG Phone 2, Blu G90 Pro, Google Pixel 1, Google Pixel 3, Google Pixel 3a XL, Huawei P20, LG G3, LG G6, LG K20, LG Revolution, LG Stylo, LG Stylo 4, LG Stylo 5, LG, Tribute HD, Motorola Moto G Power, Motorola Moto G4, Motorola Moto G7 Play, Motorola Moto G7 Power, Motorola Moto X4, Nokia 5.3, Nokia 6.1, OnePlus 6, OnePlus 8T, Realme Note 6 Pro, Samsung Galaxy, Samsung Galaxy A10, Samsung Galaxy A10E, Samsung Galaxy A20, Samsung Galaxy A71, Samsung Galaxy J3, Samsung Galaxy J3V, Samsung Galaxy J7, Samsung Galaxy Note 10+, Samsung Galaxy Note 20 Ultra, Samsung Galaxy S10, Samsung Galaxy S10+, Samsung Galaxy S10E, Samsung Galaxy S20, Samsung Galaxy S20 FE, Samsung Galaxy S20 Ultra, Samsung Galaxy S6, Samsung Galaxy S7, Samsung Galaxy S7E, Samsung Galaxy S8, Samsung Galaxy S8+, Samsung Galaxy S9, Samsung Galaxy S9+, Samsung Galaxy S20+, Techno Camon 15

**Supplementary Table 4.** Table of Sentence Verification statement stimuli

| True | | False | |
|---|---|---|---|
| **Non-negated** | **Negated** | **Non-negated** | **Negated** |
| Elephants are large. | Elephants are not small. | Elephants are small. | Elephants are not large. |
| Cars have tires. | Cars do not have wings. | Cars have wings. | Cars do not have tires. |
| Grass is green. | Grass is not blue. | Grass is blue. | Grass is not green. |
| Ice is cold. | Ice is not warm. | Ice is warm. | Ice is not cold. |
| Boulders are heavy. | Boulders are not light. | Boulders are light. | Boulders are not heavy. |
| Rocks are hard. | Rocks are not soft. | Rocks are soft. | Rocks are not hard. |
| Dogs bark. | Dogs do not meow. | Dogs meow. | Dogs do not bark. |
| Apples are fruit. | Apples are not vegetables. | Apples are vegetables. | Apples are not fruits. |
| Candy is sweet. | Candy is not salty. | Candy is salty. | Candy is not sweet. |
| Knives are sharp. | Knives are not blunt. | Knives are blunt. | Knives are not sharp. |
| Villains are evil. | Villains are not kind. | Villains are kind. | Villains are not evil. |
| Fire is hot. | Fire is not cold. | Fire is cold. | Fire is not hot. |
| The sun is bright. | The sun is not dim. | The sun is dim. | The sun is not bright. |
| Rockets are fast. | Rockets are not slow. | Rockets are slow. | Rockets are not fast. |
| Car horns are loud. | Car horns are not quiet. | Car horns are quiet. | Car horns are not loud. |
| Turtles are slow. | Turtles are not fast. | Turtles are fast. | Turtles are not slow. |
| Giraffes are tall. | Giraffes are not short. | Giraffes are short. | Giraffes are not tall. |
| Garbage smells bad. | Garbage does not smell good. | Garbage smells good. | Garbage does not smell bad. |
| Heroes are helpful. | Heroes are not useless. | Heroes are useless. | Heroes are not helpful. |
| Kids like to play. | Kids do not like to do homework. | Kids like to do homework. | Kids do not like to play. |
| Diamonds are shiny. | Diamonds are not dull. | Diamonds are dull. | Diamonds are not shiny. |

**Supplementary Table 5.** Table of non-mutually exclusive errors constituting trial removal by device

| Error Type | Percent Total Trials | | |
|---|---|---|---|
| | Computer | Tablet | Smartphone |
| Recording error | 0.7 (0 - 22.6) | 4.5 (0 - 96.8) | 4.3 (0 - 92.9) |
| Insufficient data points | 1.7 (0 - 59.1) | 7.1 (0 - 90.1) | 6.2 (0 - 86.1) |
| Reaction times > 0.1s | 0.5 (0 - 6.8) | 0.8 (0 - 18.3) | 0.7 (0 - 7.1) |
| Movement times > 3SD above participant's mean | 1.8 (0.4 - 3.2) | 1.6 (0 - 2.8) | 1.6 (0 - 3.2) |
| Reaction times > 3SD above participant's mean | 1.5 (0 - 4.4) | 1.6 (0 - 3.2) | 1.4 (0 - 3.9) |
| Incorrect trial responses | SC: 2.5 (0 - 21.4)<br>SV: 3.3 (0 - 28.6) | SC: 1.4 (0 - 46.4)<br>SV: 4.8 (0 - 47.6) | SC: 1.2 (0 - 11.9)<br>SV: 5.7 (0 - 34.6) |

**Supplementary Material References:**


Chapman, C.S., Gallivan, J.P., Wood, D.W., Milne, J.L., Culham, J.C., & Goodale, M.A. (2010b). Reaching for the unknown: Multiple target encoding and real-time decision-making in a rapid reach task. Cognition, 116(2), 168-176.

Dehaene, S., & Mehler, J. (1992). Cross-linguistic regularities in the frequency of number words. Cognition, 43(1), 1–29. https://doi.org/10.1016/0010-0277(92)90030-l

Gallivan, J. P., & Chapman, C. S. (2014). Three-dimensional reach trajectories as a probe of real-time decision-making between multiple competing targets. Frontiers in Neuroscience, 8, 215. https://doi.org/10.3389/fnins.2014.00215

Moyer, R. S., & Landauer, T. K. (1967). Time required for judgements of numerical inequality. Nature, 215(5109), 1519–1520. https://doi.org/10.1038/2151519a0

Santens, S., Goossens, S., & Verguts, T. (2011). Distance in motion: Response trajectories reveal the dynamics of number comparison. PLoS ONE, 6(9), Article e25429. https://doi.org/10.1371/journal.pone.0025429

Schwarz, W., & Ischebeck, A. (2003). On the relative speed account of number-size interference in comparative judgments of numerals. Journal of Experimental Psychology: Human Perception and Performance, 29(3), 507–522. https://doi.org/10.1037/0096- 1523.29.3.507
